# Parental-fetal interplay of immune genes leads to intrauterine growth restriction

**DOI:** 10.1101/2021.03.26.437292

**Authors:** Gurman Kaur, Caroline B. M. Porter, Orr Ashenberg, Jack Lee, Samantha J. Riesenfeld, Matan Hofree, Maria Aggelakopoulou, Ayshwarya Subramanian, Subita Balaram Kuttikkatte, Kathrine E. Attfield, Christiane A. E. Desel, Jessica L. Davies, Hayley G. Evans, Inbal Avraham-Davidi, Lan T. Nguyen, Danielle A. Dionne, Anna E. Neumann, Lise Torp Jensen, Thomas R. Barber, Elizabeth Soilleux, Mary Carrington, Gil McVean, Orit Rozenblatt-Rosen, Aviv Regev, Lars Fugger

## Abstract

Intrauterine growth restriction (IUGR) of fetuses affects 5-10% of pregnancies and is associated with perinatal morbidity, mortality and long-term health issues. Understanding genetic predisposition to IUGR is challenging, owing to extensive gene polymorphisms, linkage disequilibrium, and maternal and paternal influence. Here, we demonstrate that the inhibitory receptor, KIR2DL1, expressed on maternal uterine natural killer (uNK) cells, in interaction with the paternally-inherited HLA-C*05, an HLA-C group 2 allotype, expressed on fetal trophoblast cells, causes IUGR in a humanised mouse model. Micro-CT imaging of the uteroplacental vasculature revealed reduced uterine spiral artery diameter and increased segment length, increasing fetal blood flow resistance. Single cell RNA-Seq from the maternal-fetal interface highlighted expression programs activated by KIR2DL1-induced IUGR in several placental cell types, including degradation of extracellular matrix components, angiogenesis, and uNK cell communication, suggesting a complex response underlying IUGR. As current IUGR treatments are insufficient, our findings provide important insight for drug development.

## INTRODUCTION

Intrauterine growth restriction (IUGR) refers to pathologically reduced fetal growth and occurs in about 5-10% of pregnancies in developed countries, with higher proportions in developing countries (de Onis et al., 1998; Imdad and Bhutta, 2013; Romo et al., 2009). IUGR is one of the largest contributors to perinatal mortality and morbidity (Bernstein et al., 2000; Ego et al., 2013; Gardosi et al., 2005; Moraitis et al., 2014; Resnik, 2002; Rosenberg, 2008), and predisposes later in life to heart disease, hypertension, type 2 diabetes and stroke (Forsen et al., 2000; Frankel et al., 1996; Horikoshi et al., 2016; Leon et al., 1998; Rich-Edwards et al., 1997; Tyrrell et al., 2013; Warrington et al., 2019). IUGR can occur alone or with pre-eclampsia, a serious and systemic pregnancy complication characterised by maternal new-onset hypertension and proteinuria (Burton and Jauniaux, 2018; Steegers et al., 2010).

Despite their serious clinical implications, the causes, disease mechanisms and relationship of IUGR and pre-eclampsia are poorly understood, and effective prevention and treatment strategies are lacking (Figueras and Gardosi, 2011; McCowan et al., 2018; Nardozza et al., 2017). This is partly because studying IUGR and pre-eclampsia in human pregnancies is logistically challenging, due to the inaccessibility of tissue during critical early stages of pregnancy. Thus, there is an unmet need to better understand disease pathogenesis to identify diagnostic and therapeutic tools that improve maternal and fetal health, and long-term health of the child.

IUGR risk factors can be fetal, maternal or placental in origin. Pathogenesis is suspected to occur early in placentation when the definitive placenta is forming, and lie in placental insufficiency and dysfunction, such as higher uterine resistance to blood flow (Burton and Jauniaux, 2018; Li et al., 2018; Malhotra et al., 2019; Nardozza et al., 2017). Genetic association studies in humans suggest that uterine natural killer (uNK) cell surface receptors and fetal trophoblast ligands play a role in controlling placentation. uNK cells express a combination of inhibitory and activating killer cell immunoglobulin-like receptors (KIRs) on their cell surface, are the most abundant lymphocytes at the maternal-fetal interface, especially during early to mid-gestation, and are implicated in placental vascular remodeling (Ashkar et al., 2003; Ashkar et al., 2000; Bashirova et al., 2006; Hanna et al., 2006; Hiby et al., 2010; Lima et al., 2012; Sojka et al., 2018). HLA-C alleles are KIR ligands and can modulate NK cell function. HLA-C is the only polymorphic HLA molecule expressed on the surface of fetal extravillous trophoblasts (cells derived from the outer layer of the blastocyst, defining the boundary between the mother and the fetus) and there is direct contact between uNK and trophoblast cells during placentation (Apps et al., 2009; King et al., 2000; Lanier, 2005; Moffett and Loke, 2006).

Pregnant women with two copies of the KIR A haplotype (*i.e.,* KIR AA), composed of inhibitory KIR genes, have a higher risk (odds ratio 1.32-1.93) of pregnancy complications such as IUGR, pre-eclampsia and recurrent miscarriage, especially if the fetus inherits a group 2 HLA-C allele from the father (odds ratio 2.02). In contrast, the KIR B haplotype, mostly composed of activating KIR genes, is associated with protection against pregnancy disorders and high birth weight (Hiby et al., 2014; Hiby et al., 2010; Hiby et al., 2004; Nakimuli et al., 2015). KIR and HLA genes are highly polymorphic and inherited in complex haplotypes (Bashirova et al., 2006; Lanier, 2005; Parham, 2005), making it genetically challenging to ascertain the role of specific genes in disease pathogenesis. A typical KIR A haplotype consists of seven KIR genes and two pseudogenes, with extensive allelic variation present at several of these genes. Due to its strict binding specificity to group 2 HLA-C alleles, KIR2DL1 represents a candidate gene likely to confer the genetic risk observed in pregnancy complications (Bashirova et al., 2006; Hiby et al., 2004). Nevertheless, the KIR haplotypes have extensive gene polymorphism and strong linkage disequilibrium, so determining the role of specific genes requires teasing their effects apart from the haplotype by testing with functional assays.

Recent studies have used single cell RNA-seq (scRNA-seq) to begin to shed light on the complex cellular composition of the maternal fetal interface. In particular profiling of total immune (CD45+) cells from healthy human pregnancies (Vento-Tormo et al., 2018), highlighted three major subsets of NK cells in first trimester deciduas from healthy pregnant women, all of which expressed tissue-resident markers, but differed in their immunomodulatory profiles. Another study profiled cells from healthy and early pre-eclamptic placentas at the point of delivery (Tsang et al., 2017), but did not specifically highlight populations of NK cells. By necessity, however, such studies could only explore either first trimester placentas and be focused on likely healthy pregnancies, rather than the context of IUGR and relevant defined genotypes, or study placentas at the time of delivery by which point the cellular composition of the tissue is significantly different from that of early/mid-gestation.

Here, we developed a novel humanised mouse model system in which the interaction between the inhibitory KIR2DL1 receptor on maternal uNK cells and paternally-derived HLA-C*05, an HLA-C group 2 allotype, on fetal trophoblast cells leads to IUGR in fetuses. Our system highlights the involvement of this specific KIR-HLA-C interaction in disease pathogenesis, allowing us to study its role at the maternal-fetal interface in pregnancy. We show that this interaction leads to an increase in uterine vascular resistance, evidenced by a reduction in uterine spiral artery radius and an increase in arterial segment length. We performed single cell RNA-seq (scRNA-seq) on sorted populations of uNK cells from healthy and IUGR pregnancies to build a uNK cell atlas from the maternal-fetal interface at mid-gestation, revealing seven functionally heterogenous uNK subtypes. Topic modeling with Latent Direchelet Allocation (LDA) highlighted gene programs that are significantly perturbed in IUGR, including a program with *Gzmd*, *Gzme* and *Gzmg* expression in tissue-resident uNK cells, and a program involved in lymphocyte activation in circulating uNK cells. Further scRNA-seq of unsorted cells at the maternal-fetal interface at mid-gestation revealed expression changes in the IUGR phenotype for several non-NK cell types and a change in cellular communication between these cell types and uNK cells, suggesting a coordinated response leads to IUGR manifestation in pregnancy.

## RESULTS

### A humanised HLA-C*05 and KIR2DL1 transgenic mouse model

To characterise the role of specific KIR and HLA-C genes at the maternal-fetal interface, we used KIR2DL1 from the KIR A haplotype due to its strict specificity to bind group 2 HLA-C alleles expressed on fetal trophoblast cells. We chose *KIR2DL1*0030201* because it is a common KIR2DL1 inhibitory allele typically found on the KIR A haplotype. Correspondingly, we chose HLA-C*05 (*HLA-C*05:01:01:01*), a common group 2 HLA-C allele. We developed humanised single transgenic mice with physiological expression of HLA-C*05 or KIR2DL1 and double transgenic mice expressing both HLA-C*05 and KIR2DL1.

The HLA-C*05 transgenic mice expressed HLA-C*05 on all MHC-class I expressing cells as HLA-C*05 was driven by the endogenous *HLA-C*05* promoter. To facilitate interaction of the HLA-class I molecule with murine CD8 (and hence the T cell receptor), we replaced the *α*3 domain of HLA-C*05 with its murine counterpart from H2-K^b^, along with the adjacent transmembrane and cytoplasmic domains—as has been done for other HLA-Class I genes (Borenstein et al., 2000) (**Fig. 1a**). We confirmed that HLA-C*05 was expressed in all cells of all tested organs using an anti-HLA-C antibody and qPCR primers specific for HLA-C (**Fig. 1b and S1a**). We further confirmed that different populations of immune cells in the spleen or thymus, as well as thymic epithelial cells (TECs), expressed HLA-C*05 (**Fig. 1c,d**), demonstrating physiological expression of the transgene in the mouse model. HLA-C*05-expressing splenocytes responded to *in vitro* stimuli such as IFN*γ* or LPS by upregulating HLA-C expression (**Fig. 1e,f and S1b, c**), in agreement with previous observations (Otsuka et al., 1991).

**Fig. 1.**
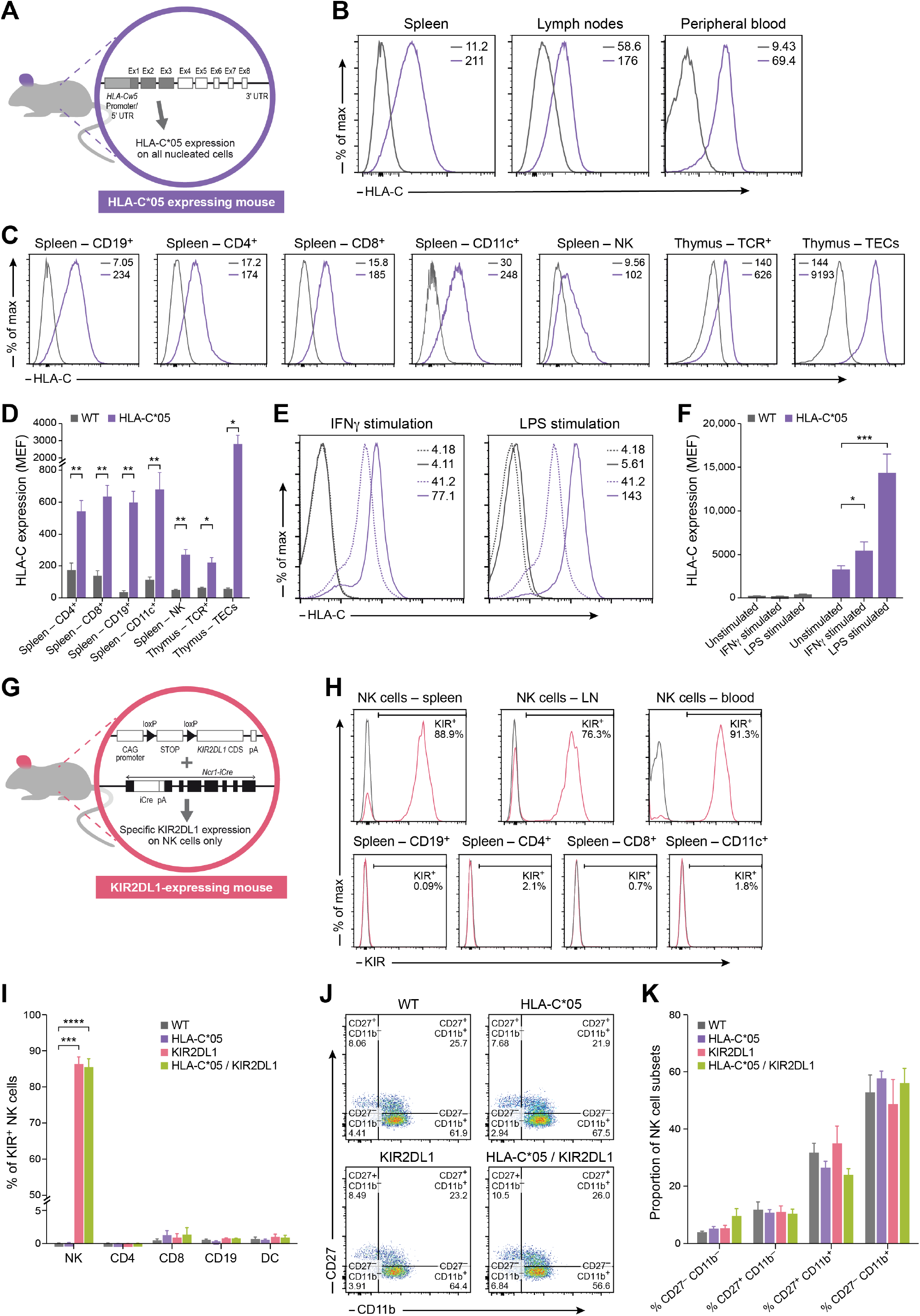
Development and characterisation of the HLA-C*05 and KIR2DL1 transgenic mouse model. **(a)** Schematic of construct used to make HLA-C*05 transgenic mice. The grey shaded region is from the *HLA-C* allele, and the white shaded region is from the murine *H-2K^b^* allele. **(b and c)** Representative cell surface expression of HLA-C*05 on total cells from organs **(b)** and on gated cell subsets from the spleen (**C**). NK cells are NKp46^+^CD3^−^TCR^−^ cells, thymus TCR+ cells are CD3^+^TCR^+^ and TECs are Epcam^+^CD45^−^ cells. Numbers denote mean florescence intensity (MFI). **(d)** HLA-C cell surface expression plotted as molecules of equivalent fluorochrome (MEF) **(e)** Representative cell surface HLA-C*05 expression on spleen CD19^+^ cells cultured with or without stimulation. Dashed lines represent unstimulated cells and solid lines represent stimulated cells. Numbers denote MFI. **(f)** HLA-C cell surface expression plotted as MEF from CD19^+^ splenocytes **(g)** Schematic of *KIR2DL1* and *Ncr1-iCre* gene expression construct used to make NK cell-specific KIR2DL1-expressing mice. **(h)** Representative cell surface expression of KIR2DL1 on NKp46^+^CD3^−^TCR^−^ NK cells in different organs or on different cell subsets from spleen. Numbers denote percentage of KIR^+^ cells. **(i)** Percentage of KIR^+^ cells in immune cell subsets from the spleen. **(j and k)** Cell surface staining of CD27 and CD11b on gated splenic NK cells. HLA-C*05 mice are shown in purple, WT mice in grey, KIR2DL1-expressing mice in pink and HLA-C*05/KIR2DL1 double transgenic mice in green. Mean + SEM is shown. (**d**) n = 3-10 per group. (**f**) n= 4-7 per group. (**i**) n = 4-11 per group. (**k**) n= 4-10 per group. *p < 0.05, **p < 0.01, ***p < 0.001, ****p < 0.0001, Mann-Whitney test.

To model the fact that KIR genes are predominantly expressed on NK cells in humans (Bashirova et al., 2006; Vilches and Parham, 2002), we generated mice where KIR2DL1 is restricted to murine NK cells. To this end, we generated transgenic mice with a loxP flanked stop cassette between the *KIR2DL1* coding DNA sequence and CAG promoter, and mated them to mice expressing iCre-recombinase in cells expressing the NK cell marker, *Ncr1* (*i.e.*, *Ncr1-iCre* BAC transgenic mice) (Eckelhart et al., 2011) (**Fig. 1g**), to generate KIR2DL1-expressing mice. Staining immune cells using anti-KIR antibody demonstrated specific and high expression of KIR2DL1 on NK cells only, with expression in ∼75-90% of NK cells (**Fig. 1h,i**).

Expression of KIR2DL1 or HLA-C*05 did not alter the proportion of NK cells (**Fig. S1d**) or other immune cell types (**Fig. S1e,f**) in the lymphoid compartments of the transgenic mice expressing HLA-C*05, KIR2DL1 or both HLA-C*05 and KIR2DL1, and NK cell receptor and transcription factor expression essential for mouse NK cell development remained unaltered (**Fig. S1g-i**). To minimize perturbations introduced to the transgenic model system, we did not knockout any mouse MHC class I or NK cell receptor molecules or abrogated any immune cell types. We further confirmed that NK cell development and maturation was normal and comparable to that observed in wild-type (WT) littermates by testing CD27 and CD11b expression, genes that represent distinct NK-cell development stages (Chiossone et al., 2009) (**Fig. 1j,k and S1j,k**).

In summary, to test the hypothesis that KIR2DL1 and HLA-C*05 in combination play a pathogenic role in the development of IUGR, we developed a novel humanised transgenic model system with specific expression of the inhibitory KIR2DL1 receptor on murine NK cells and the corresponding expression of the KIR2DL1 ligand, HLA-C*05, on all MHC-class I expressing cells in mice.

### KIR2DL1 and HLA-C*05 are functional in the transgenic mice

We next validated that both KIR2DL1 and HLA-C*05 were functional in the transgenic setting. First, to test if signalling via the inhibitory KIR2DL1 receptor impacted NK cell responses, we stimulated splenocytes from WT, HLA-C*05- or KIR2DL1-expressing transgenic mice *in vitro* using antibodies that activate NK cell receptors (NKp46 or Ly49D), observing the expected increase in IFN*γ* production by NK cells. Presence of *α*-KIR2DL1 antibody along with *α*-NKp46 or *α*-Ly49D led to significant decrease (p-value < 0.05, Mann-Whitney test) in IFN*γ* production in KIR2DL1-expressing splenocytes, but not in HLA-C*05 or WT splenocytes (**Fig. 2a,b**), confirming inhibitory signalling via the KIR2DL1 transgene.

**Fig. 2.**
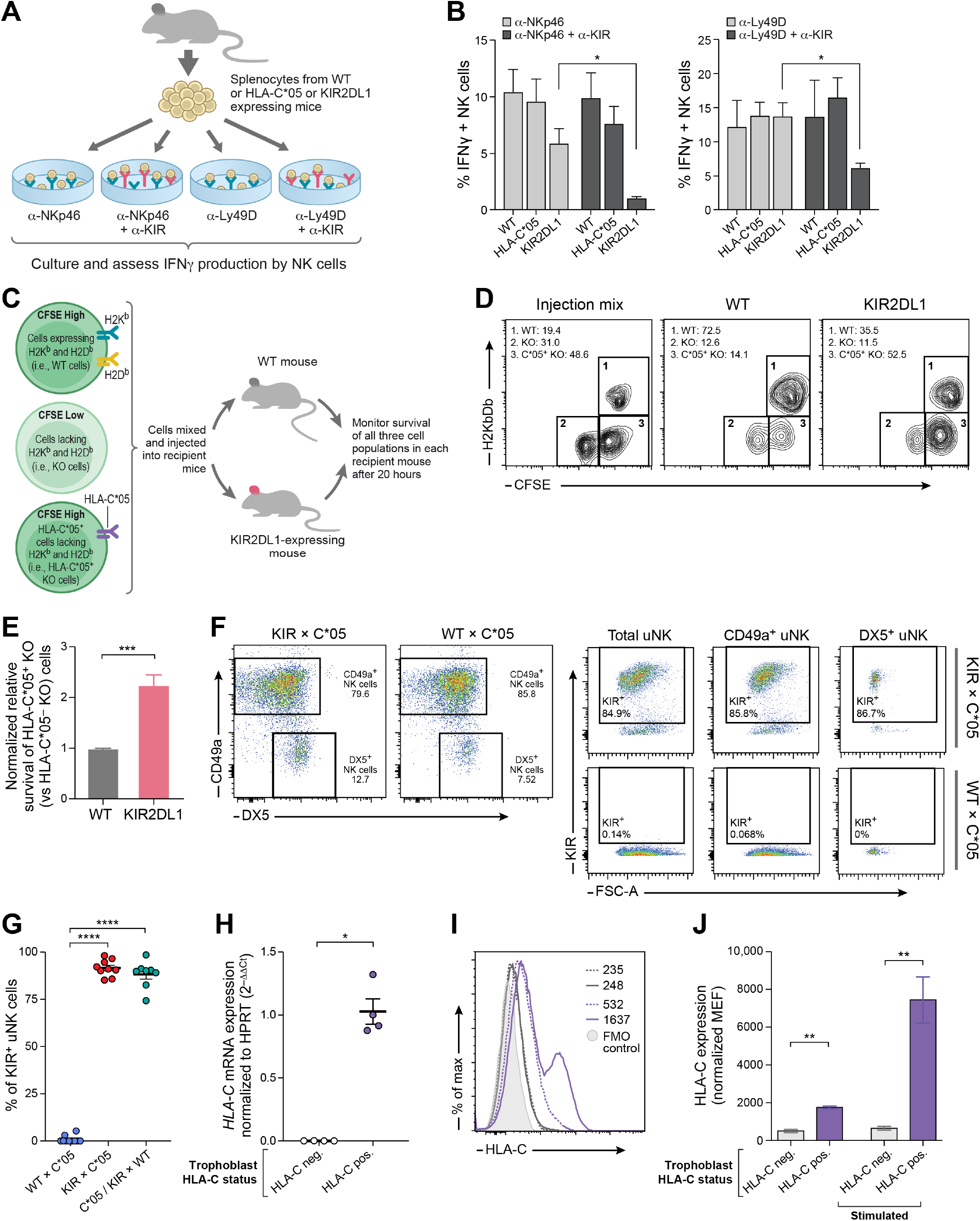
The KIR2DL1 transgene recognises HLA-C*05 and modifies NK cell effector function, and transgene expression is verified at the maternal-fetal interface. (**a**) Experimental design of NK cell stimulation experiments. (**b**) Staining for IFN*γ*^+^ NK cells (CD3^−^TCR^−^ Nkp46^+^ or CD3^−^TCR^−^DX5^+^) upon culture with antibodies depicted in panel (a). (**b**) Experimental design of adoptive transfer experiments. (**d**) Representative staining of cellular injection mix and CFSE^+^ splenocytes harvested from recipient mice. Numbers denote percentage of each gated cell population. (**e**) Relative survival of HLA-C*05^+^ KO cells compared to survival of HLA-C*05^−^KO cells (normalised to WT mice). (**f**) Representative flow cytometric staining of DX5 and CD49a on uNK cells (CD3^−^CD19^−^TCR^−^ CD45^+^NKp46^+^CD122^+^ cells) isolated from the implantation sites at gd9.5 from the different mating crosses. KIR staining is shown on total uNK cells, CD49a^+^ or DX5^+^ uNK subsets. (**g**) Percentage of KIR^+^ uNK cells in different mating crosses at gd9.5. Crosses are mentioned as female x male. (**h**) HLA-C*05 mRNA expression on fetal trophoblast cells from implantation sites at gd9.5 or gd10.5. (**i**) Representative HLA-C*05 cell surface expression on fetal trophoblast cells isolated from placenta at gd12.5, and cultured with or without stimulation. Dashed lines represent unstimulated cells and solid lines represent stimulated cells. HLA-C*05^+^ trophoblast cells are shown in purple and HLA-C*05^−^ trophoblast i.e. WT cells are shown in grey. The filled histogram represents fluorescence minus one (FMO) control. Numbers denote MFI. (**j**) HLA-C cell surface expression plotted as MEF normalised to FMO controls. Mean + SEM is shown. (**b**) n = 4-9 per group for *α*-Ly49D stimulation, n= 3-10 per group for *α*-NKp46 stimulation. (**e**) n= 6-9 per group. (**g**) n= 8-10 per group. (**h**) n = 4 per group. (**j**) n = 5 per group. *p < 0.05, **p < 0.01, ***p < 0.001, ****p < 0.0001, Mann-Whitney test.

We also assessed KIR2DL1 binding by HLA-C*05 *in vitro* by incubating the KIR2DL1-expressing NK cell line (YT-KIR2DL1) with 721.221-C*05 transfectants and quantifying intracellular IFN*γ* to measure NK cell activation. Co-culture with 721.221-C*05 transfectants, but not the control 721.221-vector or 721.221-C*07 transfectants, dampened the production of IFN*γ* by KIR2DL1-expressing NK cell line, confirming that HLA-C*05 (but not HLA-C*07 or the empty vector transduced cells) was bound by KIR2DL1 (**Fig. S1l**).

Next, to assess whether the KIR2DL1 transgene conferred inhibitory signals to NK cells upon recognition of HLA-C*05 *in vivo*, we performed adoptive transfer experiments. We isolated splenocytes from either knockout mice of murine MHC class I molecules, H2K^b^ and H2D^b^ (“KO cells”), WT mice (normal expression of H2K^b^ and′ H2D^b^, “WT cells”) or HLA-C*05 transgenic mice bred to H2K^b^ and H2D^b^ KO (“HLA-C*05^+^ KO cells”). We labelled KO, WT and HLA-C*05^+^ KO splenocytes with different concentrations of the florescent cell labelling dye CFSE to distinguish them, and injected them into either WT or KIR2DL1-expressing transgenic mice. As expected, KO cells showed increased rejection in both WT or KIR2DL1-expressing recipient mice due to missing-self recognition, given the capacity of NK cells to attack cells that do not express sufficient levels of MHC class I molecules of the host (Karre et al., 1986). The HLA-C*05^+^ KO cells were protected from NK-cell mediated rejection in KIR2DL1-expressing mice (but not WT mice) (p-value < 0.001, Mann-Whitney test) (**Fig. 2c-e**), showing that HLA-C*05 engagement by the KIR2DL1-expressing NK cells repressed mouse NK cell activating pathways.

### KIR2DL1 and HLA-C*05 are expressed at the maternal-fetal interface

We next evaluated the expression of the KIR2DL1 and HLA-C*05 transgenes at the maternal-fetal interface in two different mating combinations: (1) KIR2DL1-expressing female mated with a HLA-C*05 expressing male (KIR x C*05), chosen because in humans there is a higher risk of IUGR when the mother has the KIR A haplotype (always carrying KIR2DL1) and the father contributes the group-2 HLA-C allele (Hiby et al., 2014; Hiby et al., 2010; Hiby et al., 2004; Nakimuli et al., 2015); and (2) WT female mated with a HLA-C*05 expressing male (WT x C*05), as a control. HLA-C*05 mating males were homozygous for HLA-C*05, such that all fetuses in the litter would have the paternally-derived HLA-C*05 allele. Because the frequency of uNK cells peaks at early to mid-gestation and declines in late pregnancy (Lima et al., 2012; Sojka et al., 2018), we isolated uNK cells from mid-gestation (gestation day (gd) 9.5) from the maternal-fetal interface of KIR x C*05 and WT x C*05 matings.

In KIR x C*05, KIR2DL1 was expressed in ∼85-90% of all uNK cells—including CD49^+^ or DX5^+^ cell subsets, known to distinguish tissue resident (trNK) and conventional (cNK) NK cells (cNK cells circulate in the blood and spleen), respectively (Sojka et al., 2014); no KIR staining was observed in WT x C*05 (**Fig. 2f,g**). A third control, where a double transgenic female expressing HLA-C*05 and KIR2DL1 was mated with a WT male (C*05/KIR x WT), also expressed KIR in ∼85-90% of all uNK cells (**Fig. 2g**). The proportion of total uNK cells or CD49a^+^ and DX5^+^ uNK cell subsets was comparable across mating combinations (**Fig. S1m,n**).

We assessed expression of HLA-C*05 at the maternal-fetal interface in uterine tissue isolated from implantation sites at gd9.5 or gd10.5 of an HLA-C*05 negative female mated to either an HLA-C*05 positive or HLA-C*05 negative male. This meant that HLA-C*05 expression at the maternal-fetal interface could be ascribed only to the paternally-derived HLA-C*05 allele on fetal trophoblast cells. We detected HLA-C mRNA expression in C*05^+^ but not in C*05^−^ fetal cells (**Fig. 2h**), and HLA-C DNA, by genotyping of the fetus using genomic DNA isolated from embryonic tissue. Furthermore, we isolated and cultured fetal trophoblast cells from gd12.5 placentas and validated the paternally-derived HLA-C expression in HLA-C*05^+^ trophoblast cells by flow cytometry. Cultured fetal trophoblast cells were predominantly CD45^−^Cytokeratin-7^+^ (data not shown), and *in vitro* stimulation with LPS + IFN*γ*led to a significant upregulation of HLA-C expression in HLA-C*05^+^ (p-value<0.01, Mann-Whitney test), but not HLA-C*05^−^, trophoblast cells (**Fig. 2i,j**) — as shown previously (King et al., 2000).

Taken together, these results demonstrated the functionality of the KIR2DL1 and HLA-C*05 transgenes in the mouse model system and their expression at the maternal-fetal interface, confirming the validity of this model to study IUGR-relevant disease mechanisms.

### Maternal KIR2DL1 and paternal HLA-C*05 expression leads to IUGR in fetuses

Because birth and fetal weights are markers for intrauterine development, we investigated the effect of the KIR-HLA-C interactions in mediating IUGR by assessing fetal weight in the three mating combinations described above (KIR x C*05, WT x C*05 and C*05/KIR x WT), as well as two additional controls (KIR x WT and WT x WT). Weight was measured just before birth, at gd18.5, to avoid milk uptake as a confounding factor.

Pregnancies involving females with KIR2DL1 expression on uNK cells and males with expression of HLA-C*05 (KIR x C*05 mating) led to IUGR in the fetuses, evidenced by a reduction in the average fetal weight of the embryos compared to all other mating combinations. The progeny from the KIR x C*05 mating were ∼12% lower in weight compared to the control WT x C*05 mating (**Fig. 3a,** p-value < 10^−3^, linear mixed-effects model) and ∼56% of KIR x C*05 fetuses were below the 10^th^ weight percentile of control WT x C*05 fetuses (with 36% below the 5^th^ percentile of control WT x C*05 fetuses) (**Fig. 3b**). The placental weight was very similar across mating combinations (**Fig. 3c**), concordant with the lack of observed lesions or pathology in the placenta assessed by H&E staining (data not shown). The observed 12% reduction in average fetal weight and the skewed fetal weight distribution of the KIR x C*05 fetuses mirrors clinical observations (Figueras and Gardosi, 2011; Hiby et al., 2014; Moraitis et al., 2014).

**Fig. 3.**
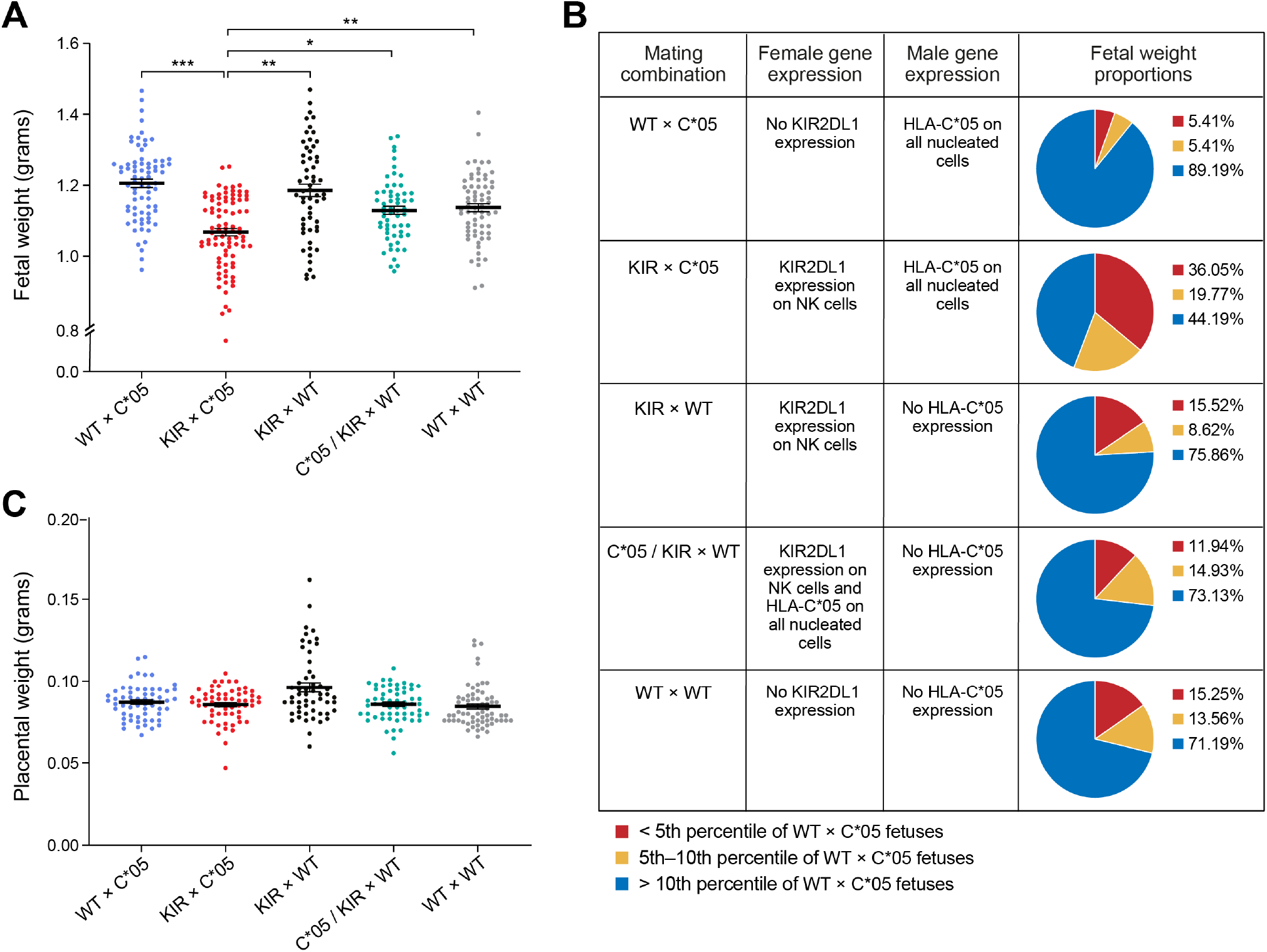
Expression of maternal KIR2DL1 and paternal HLA-C*05 leads to intrauterine growth restriction in fetuses. (**a**) Fetal weight and (**c**) placental weight determined at gd18.5 of progeny from different mating combinations involving WT, KIR2DL1-expressing and HLA-C*05 transgenic mice. Crosses are mentioned as female x male. Mean ± SEM is shown. n = 8-12 litters per group. *p < 0.05, **p < 0.01, ***p < 0.001, linear mixed effects model. **(b**) Table showing distribution of fetal weight proportions from each mating combination compared to the WT x C*05 control mating. Numbers depict percentages of fetuses whose weight was below the 5^th^ percentile, between 5^th^ – 10^th^ percentile and above the 10^th^ percentile of the WT x C*05 mating controls.

Pregnant KIR2DL1-expressing females from the KIR x C*05 mating did not show evidence of hypertension throughout the course of gestation and post-partum (**Fig. S2a**) or of proteinuria (**Fig. S2b**), both hallmarks of pre-eclampsia (Steegers et al., 2010). Increased levels of the anti-angiogenic molecule soluble fms-like tyrosine kinase 1 (sFLT-1) and reduced levels of placental growth factor (PIGF) are associated with an increased risk of developing pre-eclampsia in suspected pregnancies (Duhig et al., 2019; Levine et al., 2004; Zeisler et al., 2016), however, their usability in prediction of IUGR is limited (Conde-Agudelo et al., 2013). We did not observe any change in the circulating levels of sFLT-1, soluble Endoglin, or PIGF in the plasma of pregnant females from the KIR x C*05 mating combination (**Fig. S2c-e**).

Collectively, these results demonstrate the impact of specific KIR and HLA genes on IUGR, with direct evidence of the involvement of KIR2DL1 and HLA-C*05 genes. Furthermore, they corroborate and elaborate on human genetic association studies showing that there was a significant increase in fetal growth restriction, pre-eclampsia and recurrent miscarriage in pregnancies in women with two copies of the KIR A haplotype, but only when there was either a paternally-inherited group 2 HLA-C gene in the fetus, or when the fetus had more group 2 HLA-C genes than the mother (Hiby et al., 2010). Moreover, the impact we observed on IUGR in the fetuses was not associated with any manifestations of a systemic disease such as pre-eclampsia. This could suggest that that additional risk factors or mechanisms are required for pre-eclampsia manifestation.

### Maternal KIR2DL1 and paternal HLA-C*05 expression leads to changes in uterine spiral arteries during gestation

Because compromised uterine circulation may contribute to IUGR (Burton and Jauniaux, 2018), we looked for changes in maternal uterine spiral arteries that feed the developing fetus in the KIR x C*05 mice at gd10.5, a point when coiling spiral arteries in the decidua have formed (Adamson et al., 2002). Mice were perfused with an X-ray opaque contrast agent into the uteroplacental circulation and imaged using micro-computed tomography (micro-CT) to preserve the *in vivo* morphology of the circulatory spaces (**Fig. 4a**). We manually segmented the spiral arteries in both the KIR x C*05 (IUGR) and WT x C*05 (control) mating combinations (**Fig. 4b,c**) and skeletonized them with homotopic thinning (**Fig. S3a,b**).

**Fig. 4.**
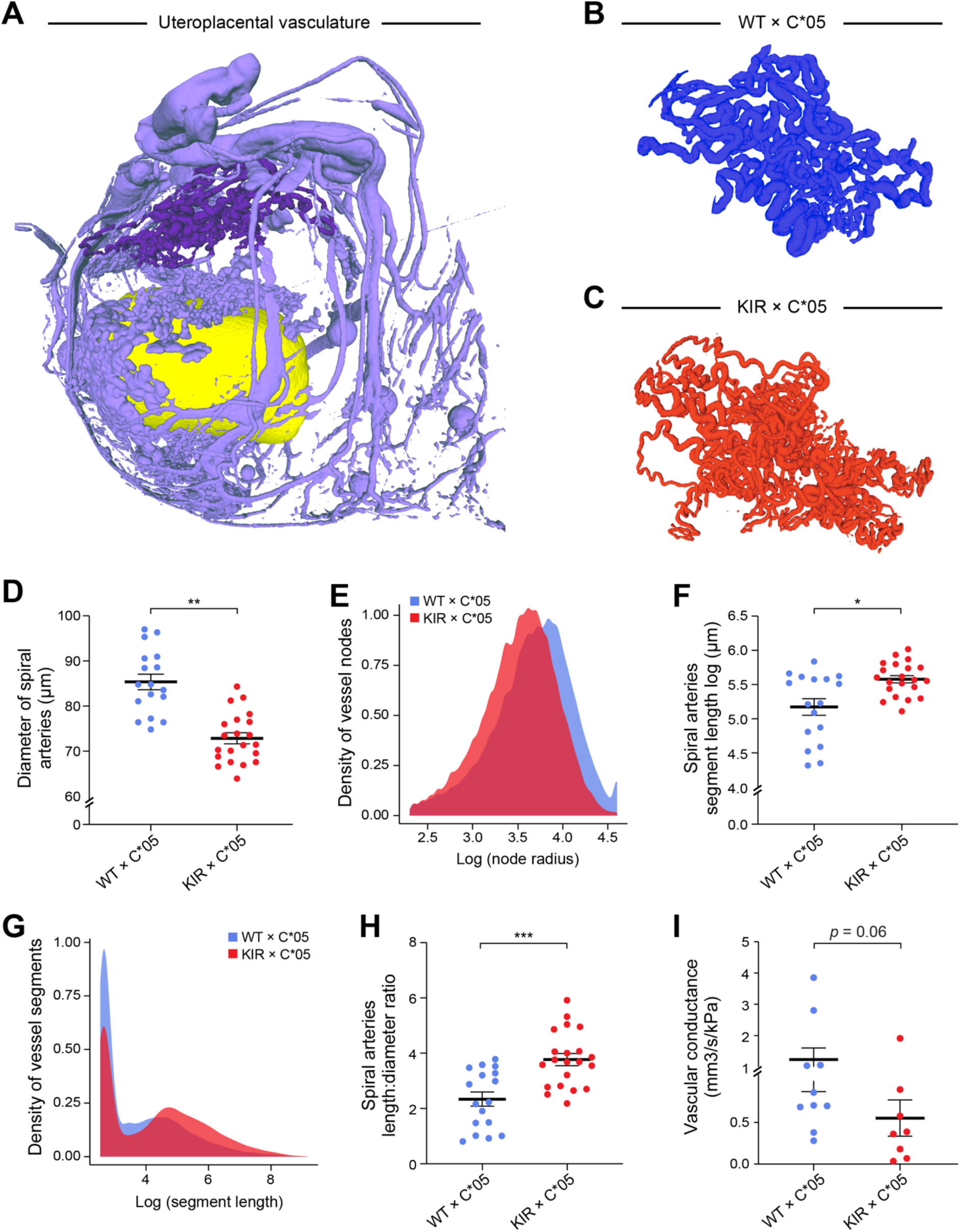
Combination of maternal KIR2DL1 and paternal HLA-C*05 leads to changes in uterine spiral arteries during gestation. (**a**) Representative uteroplacental circulation from WT x C*05 mating at gd10.5 visualised following perfusion with an X-ray contrast agent and imaging by micro-CT. Spiral arteries are shown in dark purple and the yolk sac is shown in yellow. (**b and c**) Representative spiral artery vasculature images from the different mating crosses at gd10.5. (**d and f**) Spiral artery diameter (**d**) or segment length (**f**) is shown. (**e and g**) Radii distribution of spiral artery vessel nodes (**e**) and distribution of segment lengths (**g**) is shown. (**h**) Ratio of the mean of the spiral artery segment length and mean of the spiral artery diameter of each implantation site is shown. (**i**) Total network fluid conductance based on the Poiseuille flow model. Mean ± SEM is shown. (**d, e, f, g and h**) n = 17–21 implantation sites per group. (**i**) n= 8-10 implantation sites per group. *p < 0.05, **p < 0.01, ***p < 0.001. Linear mixed effects model used for (d) and (f). Mann-Whitney test used for (h) and (i).

The mean diameter of the spiral arteries was reduced by ∼15% in the KIR x C*05 IUGR mice *vs*. WT x C*05 control (p-value < 0.01, linear mixed effects model, **Fig. 4d**), also evident by a shift in radii distribution of the spiral artery vessel nodes (**Fig. 4e and S3c**). Furthermore, a ∼38% increase in the mean spiral artery segment length was observed in KIR x C*05 IUGR mice compared to WT x C*05 controls (**Fig. 4f**, p-value < 0.05, linear mixed effects model), with a skew in the spiral artery segment length distributions in the opposite direction of that observed for the radii (**Fig. 4g and S3d**). There was a trend in the KIR x C*05 IUGR mice to have fewer spiral artery vessel segments (p=0.07, Mann-Whitney test), while the total length of spiral artery vasculature and the segment volume remained unaltered (**Fig. S3e-g**).

Due to the skewed spiral artery length-to-diameter ratio in KIR x C*05 mice (**Fig. 4h**), we also quantified the total spiral artery network resistance by imposing a pressure gradient between the inlets and outlets and calculating the total outflow. The vascular conductance through the IUGR KIR x C*05 spiral artery network tended to be lower than in WT x C*05 mating controls (p=0.06, Mann-Whitney test) (**Fig. 4i**), suggesting a higher resistance to blood flow through the spiral artery network. Collectively, these findings demonstrated that the IUGR phenotype observed in KIR x C*05 matings was associated with inadequate placentation and a change in remodeling of uterine spiral arteries during gestation.

### Atlas of uNK cells at the mid-gestation mouse maternal-fetal interface

To better understand uNK function and diversity at the maternal-fetal interface, we next built a single cell atlas of NK cells, using both full length scRNA-seq (with SMART-Seq2 (Picelli et al., 2014; Trombetta et al., 2014), providing deeper coverage per cell, **Methods**), and 3’ droplet-based scRNA-Seq (profiling large cell numbers, **Methods**), profiling isolated uNK cells at gd9.5 from the three different mating groups: KIR x C*05 (IUGR), WT x C*05 (control 1) and C*05/KIR x WT mice (control 2) (**Fig. 5a**). For full length scRNA-seq, we first FACS sorted uNK cells (NCR1^+^CD3ε^−^TCRb^−^ CD19^−^CD45^+^CD122^+^) into either trNK (CD49a^+^) or cNK (DX5^+^) cells, collecting 1,091 and 1,085 cell profiles, respectively, across five mice each for the three mating groups (**Fig. 5b, S4a-c**). For droplet based scRNA-seq, we used total uNK cells, profiling 30,147 high quality uNK cells across 3-4 mice from each of the three mating groups (**Fig. 5d, S4d-f**). Each mouse represents cells pooled from the maternal-fetal interface of multiple implantation sites within the same litter.

**Fig. 5.**
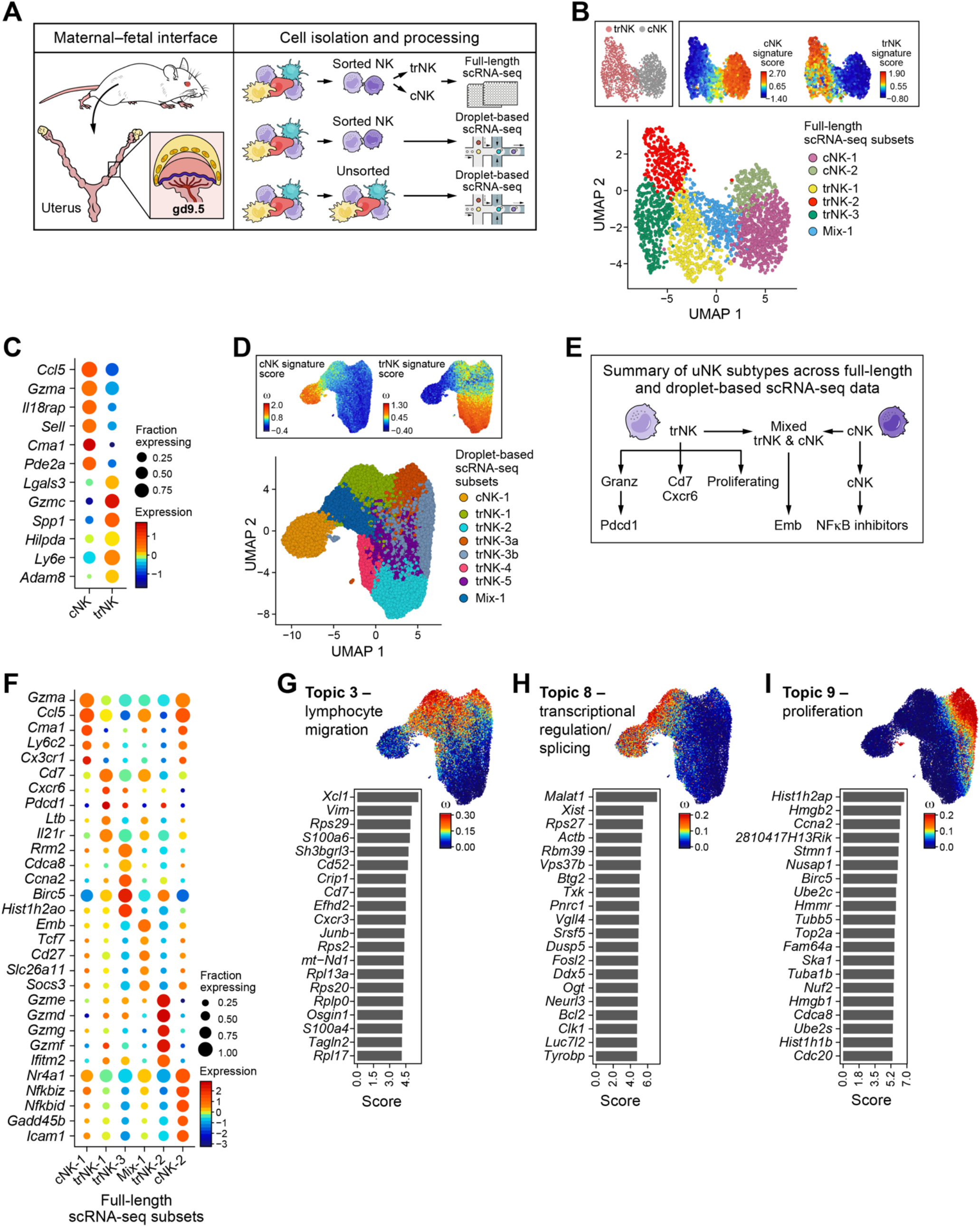
Heterogeneity of NK cells at the mouse maternal-fetal interface. (**a**) Overview of study design. (**b**) UMAP embedding of full length scRNA-seq on sorted NK cells colored by subtype as cNK or trNK (determined by FACS sorting and verified using the cNK and trNK signature score) or by Louvain cluster label. (**c**) Dot plot showing top six cNK and trNK marker genes. (**d**) UMAP embedding of NK cells profiled by droplet-based scRNA-seq and colored by Louvain cluster or the cNK and trNK signature score. (**e**) Summary of NK subtypes identified in sorted uNK cells derived from both full length and droplet based scRNA-seq data. (**F**) Dot plot of top five marker genes for each cluster shown in panel **(b)**. (**g-i**) Left: top twenty genes driving the topic; the ‘score’ has been scaled to improve visualization. Right: UMAP embedding of droplet based scRNA-seq uNK data (same as in panel **d**) colored by the weight of each cell in the topic. Topic 3 (G), Topic 8 (H), Topic 9 (I). n= 2176 cells for full length and 30,147 cells for droplet-based NK data. In all plots, cells from all three genotypes (WT x C*05, KIR x C*05, C*05/KIR x WT) are included. In dot plots, the size of the dot represents the fraction of cells with nonzero expression of each gene, and the color of the dot represents the average nonzero gene expression.

Based on the full length scRNA-seq, cNK and trNK cells formed two distinct subsets in a low dimensionality embedding (**Fig. 5b**), and could be distinguished by signatures of differentially expressed genes as determined by Mann Whitney test (**Fig. 5b,c**). cNKs expressed cellular migration and homing markers (*e.g.*, *Ccl5*, *Sell*), while trNKs expressed cell-cell and cell-matrix interaction markers (*e.g.*, *Adam8*, *Spp1*). The two NK populations also had distinct granzyme profiles, with cNKs and trNKs showing higher *Gzma* and *Gzmc* expression, respectively (**Fig. 5c**). trNKs had more genes detected per cell (median number of genes per cell in trNKs = 4,258 and in cNKs = 2,165; p value = 6.96 x 10^−264^, Mann Whitney U test) and a larger size by FACS (data not shown) than cNKs, consistent with higher transcriptional activity (**Fig. S4a**); uNK cell subset size differences have been demonstrated microscopically (Paffaro et al., 2003). Scoring the droplet-based profiles with the full-length based cNK and trNK signatures similarly distinguished the two populations in the low dimensionality embedding (**Fig. 5d**). As reported previously (Doisne et al., 2015), cNK cells represented a smaller proportion of total profiled uNK cells (**Fig. 5d and S1n**).

### Seven uNK cell subsets in the maternal-fetal interface

Unsupervised clustering identified six NK subsets in the full length scRNA-Seq data (**Fig. 5b**) and eight NK subsets in the droplet-based scRNA-Seq data (**Fig. 5d**); after evaluating and comparing subsets across both datasets, we annotated seven distinct NK cell subsets (concordant between the two datasets) at the maternal-fetal interface at gd9.5 (**Fig. 5e, Fig. S5a,b**). Due to the deeper gene coverage, we prioritised the subset-specific differentially expressed genes from full length scRNA-seq to annotate the six NK subtypes found in the full length data (**Fig. 5f**): (1 and 2) two cNK subtypes (cNK-1 and cNK-2) that shared several cNK marker genes, however, only cNK-2 had NF-kB inhibitors (*Nfkbiz*, *Nfkbid*, *Nkfkia*), (3) a trNK cell subset (trNK-1) enriched for cytokine receptor signalling genes and marked by *Cd7*, *Cxcr6* and *Pdcd1*; (4) *Gzmd/e/g/f*-high trNKs (trNK-2) enriched for several central and protein metabolic pathways (*e.g.*, glycolysis, p=3.1×10^−5^, Fisher’s exact test), (5) proliferating trNKs (trNK-3 or trNK-3a/3b) marked by expression of genes such as *Birc5* and *Ccna2*, and a high gene proliferation signature score (**Fig. S5c**, **Table S1**), and (6) a subset including both cNK and trNK cells (mix-1), defined by expression of *Cd27*, *Cd7* and *Emb* (the gene signature for this mix-1 subset scores higher on cNKs than trNKs, **Fig. S4g-h**). Of the above six subsets, only cNK-2 did not form a distinct subset in the droplet-based data, however, scoring its signature highlights a distinguishable population within the cNK-1 cluster (**Fig. S5a**). The seventh NK subset, *Gzmd/e/g/f*-high trNKs expressing *Pdcd1* (trNK-4), was a distinct subset only in the droplet-based data. Scoring its signature identified it within the trNK-2 subset in the full length data (**Fig. S5b**), but the cells largely overlapped cells in the full length and droplet-based trNK-2 cluster and signature, and it is possible that this subset was over-split in the droplet-based data (**Fig. S5b**). The eighth subset found in the droplet-based data (trNK-5) had high expression of the chemokine *Ccl1,* but was less well defined in the droplet data (**Fig. S5b**) and not included as a major uNK cell subset in our final census (**Fig. 5b,d,e,f**).

Overall, cells from the different genotypes, KIR x C*05 (IUGR), WT x C*05 (control 1) and C*05/KIR x WT mice (control 2), had similar distributions across most of the subsets identified by clustering, suggesting that genotype did not have a dramatic effect on the overall NK cellular categories (**Fig. S4i-n**), with the exception of the *Gzmd/e/g/f*-high, *Pdcd1*-high trNK-4 cluster (in droplet-based scRNA-seq), which was enriched for cells from the KIR x C*05 (IUGR) genotype (p < 0.05, Dirichlet-multinomial regression, as described before (Smillie et al., 2019), **Fig. S4n**). Subset trNK-4, however, was not distinct from trNK-2 in the full length data and shared many marker genes with trNK-2 in both datasets.

Collectively, these results highlighted the diversity of uNK cells during mid-gestation and broadly identified seven transcriptionally distinct subsets of cNK and trNK cells present at the maternal-fetal interface in control and IUGR mice.

### Increased lymphocyte activation and decreased ECM and tissue remodelling programs in uNK cell subsets in IUGR

Given the limited changes in the proportion of broad uNK subsets between genotypes and the continuum of uNK cell subsets, we hypothesised that IUGR may have more nuanced changes in cell intrinsic gene programs within NK subsets. To uncover those, we used topic modelling with latent Dirichlet allocations (LDA) (Bielecki et al., 2018; Dey et al., 2017; Xu et al., 2019) to learn 16 gene programs (“topics”) across the droplet based scRNA-seq profiles (**Fig. S5d**, **Methods**). This approach assigns each gene and cell a weight in every topic, indicating the importance of the gene to the topic and of the topic to the cell, with the top scoring genes defining the gene program for that topic. Cells and genes can be highly weighted for more than one topic.

Topics ranged from general biological processes (*e.g.,* ribosomal biogenesis, topic 1) to regulation of NK cell function (topic 10) (**Fig. S6**), and included cNK/trNK specific topics and ones spanning cells from both subtypes. For example, Topic 3 (“lymphocyte migration”, **Fig. 5g**) was enriched for genes involved in immune cell infiltration, lymphocyte migration and cytokine response and spanned trNK-1, trNK-3a and mix-1 cells; Topic 8 (“transcriptional regulation and splicing” **Fig. 5h**) spanned cNK-1, trNK-1 and mix-1 cells, and was enriched for genes involved in transcriptional regulation, spliceosome machinery and cell differentiation; and Topic 9 (“proliferation”, **Fig. 5i**) scored highly in the proliferating cells from both trNK (trNK-3a and trNK-3b) and cNKs (**Fig. S5c**). Some topics revealed gene programs that distinguish between cells within a single cluster (**Fig. S6**).

The cell weights for topic 6 (lymphocyte activation) and 16 (ECM and tissue remodelling) were significantly and robustly different in cells from KIR x C*05 IUGR mating *vs*. control matings (**Fig. 6a-c, Methods**). The lymphocyte activation program (Topic 6) is induced in IUGR (p-value = 5.06*10^− 9^, KS statistic = −0.049 for IUGR vs. CTR1; p-value = 4.09*10^−14^, KS statistic = −0.073 for IUGR vs. CTR2; KS test) (**Fig. 6b**), whereas the ECM and tissue remodelling program (Topic 16) is repressed (p-value = 3.3*10^−3^, KS statistic = 0.028 for IUGR vs CTR1; p-value = 3.34*10^−5^, KS statistic = 0.043 for IUGR vs CTR2; KS test) (**Fig. 6c**).

**Fig. 6.**
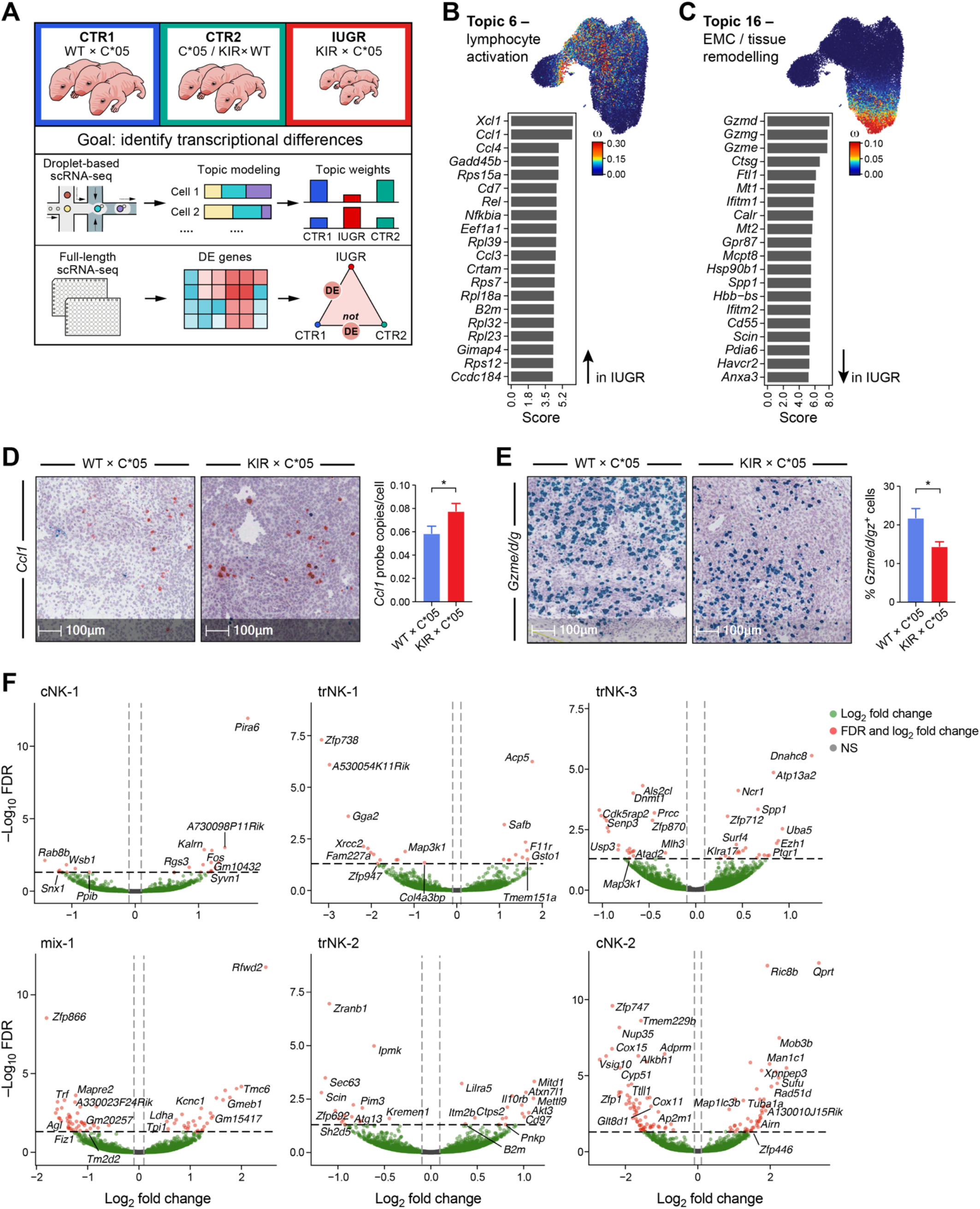
Characterisation of the IUGR phenotype in NK cells by topic modeling and differentially expressed genes. **(a)** Overview of IUGR phenotype characterisation by topic modeling and differential gene expression analysis. **(b-c)** Topic modeling of droplet-based scRNA-seq uNK cell data from the mouse maternal-fetal interface. The cell topic weight distributions for the displayed topics (Topic 6 (b), and Topic 16 (c) were significantly different between KIR x C*05 *vs.* control mating groups. Left: top 20 genes driving the topic; the ‘score’ has been scaled to improve visualization. Right, top: UMAP embedding colored by the cell weight for that topic – bright red indicates the cell is high in the topic and dark blue indicates the cell is low in the topic. **(d, e)** Re-validation of *Ccl1* (**d**) and *Gzme/d/g* (**e**) from the topics using *in-situ* hybridisation on implantation sites at gd9.5. Representative probe staining on sections from the different mating crosses is shown. Quantification depicted as percentage of positive cells or average probe copies per cell for the respective probes within a stained section is shown. **(f)** Volcano plots showing differentially expressed genes in the IUGR KIR x C*05 mating combination. Differentially expressed genes were calculated on a per cluster basis, and are stratified as such in the panel. (**d, e**) n=13-14 sections for *Ccl1* and n=15-16 sections for *Gzmd/e/g*, Mann-Whitney test.

The lymphocyte activation program induced in IUGR (Topic 6) is enriched for NF-kB signalling pathway, ribosomal genes, chemokine signalling and cytokine-cytokine receptor interaction genes (p = 0.0008, 1.37*10^−32^, 0.0131, 0.048, respectively; Fisher’s exact test). Cell weights for topic 6 are highest in the region of the droplet-based cNK-1 subset expressing NF-kB inhibitors (aligned with full length cNK-2 subset) and in the mixed cNK/trNK subset (mix-1). In the absence of proliferation, hyperactive ribosome biogenesis can indicate cellular stress (Orsolic et al., 2016). NF-kB regulates cytokine and stress-induced expression of *Gadd45b* (a topic gene involved in regulation of cell growth and apoptosis (Jin et al., 2002; Liebermann et al., 2011)), and *Cd7* is a topic gene involved in NK cell costimulatory triggering (Rabinowich et al., 1994). *In situ* hybridization on mid-gestation uterine tissue also confirmed increased expression of one of the top genes *Ccl1* in KIR x C*05 IUGR mice *vs*. controls (**Fig. 6d**).

The ECM and tissue remodelling program repressed in IUGR (Topic 16) has cell weights highest in the *Gzmd/e/g/f*-high trNK subtype (trNK-2). Top program genes (*Gzmd*, *Gzmg*, *Gzme,* but not *Gzmf*) are expressed by uNK cells in pregnancy and upregulated by IL-12 and IL-15 in uterine decidual cells (the modified uterine endometrium formed during pregnancy). These granzymes are proposed to be non-cytotoxic and play a role in ECM and tissue remodelling, as well as in parturition (Allen and Nilsen-Hamilton, 1998; Croy et al., 1997; Delgado et al., 1996). Decreased expression of *Gzmd/e/g* was confirmed by *insitu* hybridization on mid-gestation uterine tissue in KIR x C*05 IUGR mice *vs*. controls (**Fig. 6e**). Other program genes (*Ctsg*, *Mt1* and *Mt2*) play roles in tissue remodelling, cell signalling and protection against oxidative damage (Gao et al., 2018; Subramanian Vignesh and Deepe, 2017). The program gene *Havcr2* (or *Tim-3*), is associated with NK cell activation and maturation (Ndhlovu et al., 2012), and patients undergoing recurrent miscarriages have fewer Tim3^+^ NK cells—suggesting that these Tim3^+^ NK cells promote maternal-fetal tolerance (Li et al., 2017; Li et al., 2016; Subramanian Vignesh and Deepe, 2017).

We also tested for genes that are differentially expressed in cells from the IUGR phenotype within a specific NK cell subset (calculated using pseudobulk differential expression analysis, **Methods, Fig. 6a,f**). Some differentially expressed genes not highlighted by topic modeling include *Fos*, *Acp5*, *Spp1, Gsto1*, *Map3k1*, *Akt3*, and *Il10rb* (**Fig. 6f**). *Fos* encodes for one of the proteins in the AP-1 transcription complex and plays a role in the modulation of NK cell function (Marusina et al., 2008); *Acp5* encodes an enzyme (TRAP) that regulates the activity of Osteopontin (encoded by *Spp1* – also a differentially expressed gene), which has been associated with regulation of fetal growth and recurrent spontaneous abortion (Ek-Rylander et al., 1994; Fu et al., 2017); *Gsto1* is a stress response gene, which plays a role in redox homeostasis (Kodym et al., 1999); *Akt3* is a protein kinase family member, which can also promote induction of reactive oxygen species (Polytarchou et al., 2020); *Map3k1* integrates multiple signalling pathways and can also crosstalk with Wnt signalling (Meng et al., 2018); and *Il10rb* is a cell surface receptor for multiple cytokines and can stimulate activation of the JAK/STAT signaling pathway (Donnelly et al., 2004).

Together, our analyses show that in IUGR there is an increase in programs related to regulation of cell growth, apoptosis and NF-kB signalling, and decrease in programs for ECM and tissue remodelling and protection against oxidative stress.

### IUGR-associated transcriptional changes across diverse cell types at the maternal-fetal interface

To understand NK cell states in the broader context of the cellular ecosystem, we next analysed 42,869 high quality droplet-based scRNA-seq profiled from all cell types at the maternal-fetal interface at mid-gestation (gd9.5; n=3 from each mating combination) and identified transcriptional differences in IUGR by cell type (**Fig. 7a and S7a-c**).

**Fig. 7.**
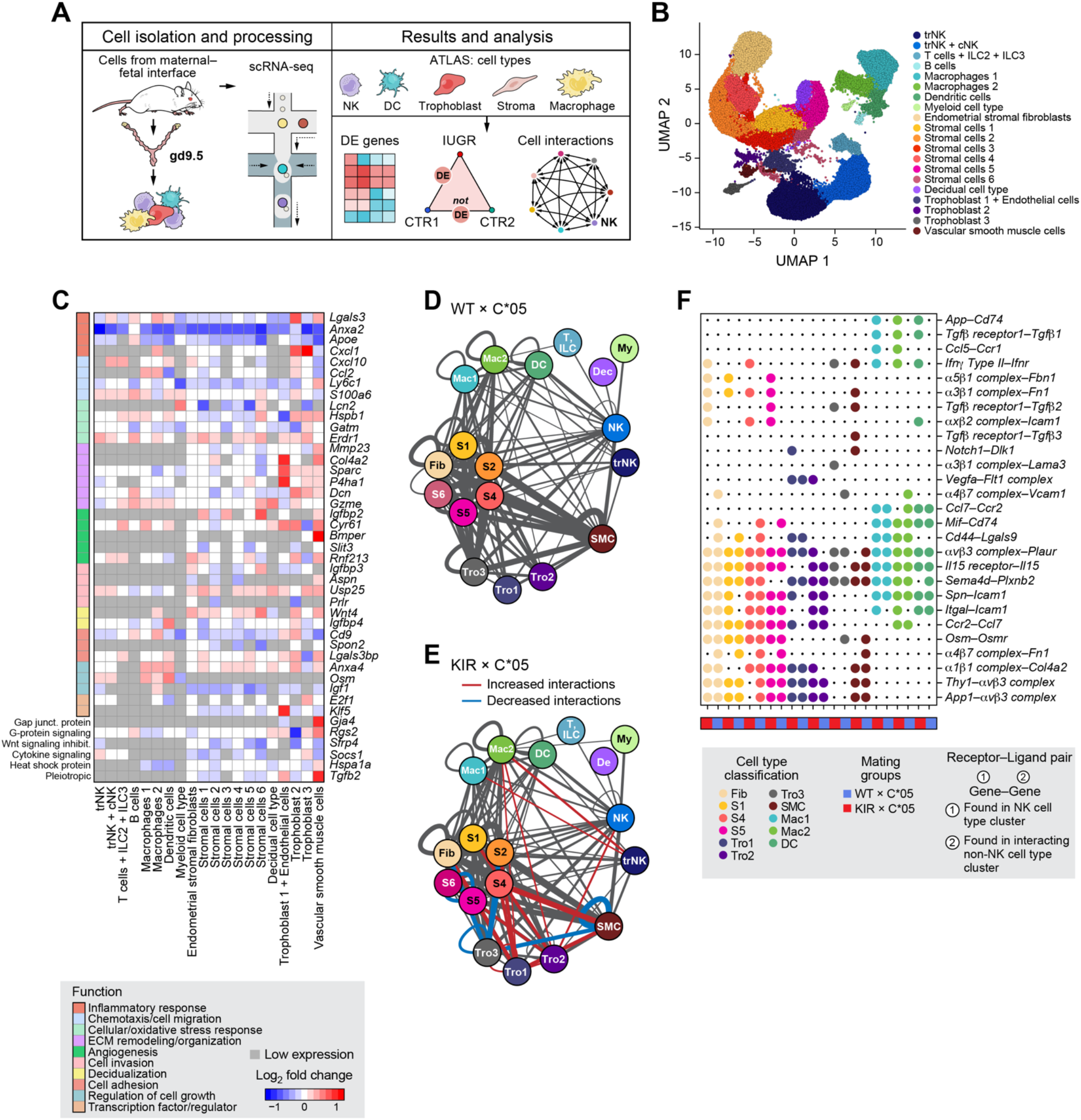
Characterisation of cell types, the IUGR phenotype and cell-cell interactions in unsorted cells from the maternal-fetal interface. **(a)** Overview of scRNA-seq analysis of unsorted cells. **(b)** UMAP embedding of unsorted cells from all three genotypes analysed using droplet-based scRNA-seq and colored by cell type classification. **(c)** Heatmap of select differentially expressed genes between KIR x C*05 and WT x C*05 mating groups. Differentially expressed genes were calculated on a per cell type basis, and every gene in this heatmap was differentially expressed in at least one cell type. The color of each square indicates the log of the average expression fold change between KIR x C*05 and WT x C*05. Bright red - higher average expression in KIR x C*05; Dark blue - lower average expression in KIR x C*05; White - no difference; grey – the gene was expressed in less than 10% in both conditions within a cell subset. The colored boxes to the left of the heatmap indicate the top functional annotation of each gene. **(d-e)** Cell-cell interaction analysis by CellPhoneDB. Cell-cell interaction network for WT x C*05 cells (**d**) and KIR x C*05 cells (**e**). The color of each circle corresponds to the cell type classification on the UMAP in panel **(b)**, and the weight of each line represents the number of significant ligand-receptor interactions found between the cell type nodes on either end, with a thicker line indicating a great number of significant interactions. Lines are only shown between cell type nodes if at least 50 significant interactions were found. Interactions increased or decreased in the KIR x C*05 group compared to the WT x C*05 group are shown in red and blue, respectively. **(f)** Select ligand-receptor pairs that were found to be unique in either KIR x C*05 or WT x C*05 groups in one of the NK cell types. The first gene listed in the label was found in one of the NK clusters, and the second gene was found in an interacting non-NK cell type cluster. A large circle indicates that the receptor-ligand pair was found between an NK cluster and the indicated cell type; cell types are colored as in panel **(b)**. For example, *App_Cd74* was found between one of the NK subtypes and macrophages 1, macrophages 2 and dendritic cells in the KIR x C*05 genotype, but not in the WT x C*05 genotype.

Unsupervised clustering and post-hoc annotation highlighted 20 cell subsets (**Methods**), spanning immune, stromal, decidual, trophoblast and vascular smooth muscle (vSMCs) cells (**Fig. 7b**), similar to findings from early human pregnancies (Vento-Tormo et al., 2018). There were nine immune cell subsets: trNKs, mixed cNKs and trNKs, (**Fig. S7d**), T cells, B cells, ILC2/3s, dendritic cells (*Cd209a*+), two macrophage subsets differing in complement (*C1qa* and *C1qb*) and MHC II (*H2-Ab1* and *H2-Eb1*) expression, and a myeloid subset (expressing *Retnlg*, *Slpi* and *S100a9* (Yang et al., 2018). There were seven stromal cell subsets: one of endometrial stromal fibroblasts (*Col3a1*, *Col1a1*, *Col1a2*, *Dcn*, *Lum, Eln* and *Postn*) and six of cell subsets at different stages of decidualisation (critical for remodeling the uterine endometrium) as reflected by the level of Wnt signalling molecules and modulators (*Wnt4*, *Wnt5a*, *Wnt6, Sfrp1*) (Ramathal et al., 2010; Zhang and Yan, 2016), TGF-*β* superfamily morphogen (*Bmp2*) (Ramathal et al., 2010; Zhang and Yan, 2016), angiogenic factors (*Angpt2* and *Angpt4)* and steroid biosynthesis genes (*Cyp11a*). One non-canonical subset was decidual-like, expressing *Prl8a2* (a decidual prolactin-related protein that contributes to pregnancy-dependent adaptations to hypoxia (Bu et al., 2016)), *Cyrab* (important in mouse embryo implantation and decidualization) and *Scgb1a1* (encoding Uteroglobin and expressed by mucosal epithelial cells (Mukherjee et al., 2007)); these cells may support decidualisation (**Fig. 7b**).

There were three trophoblast subsets: Trophoblast 1 cells expressed *H19*, which influences trophoblast cell migration and invasion (Zuckerwise et al., 2016), and other trophoblast genes, including *Bex1*, *Krt8* and *Mest*, and included cells expressing endothelial markers (*e.g.*, *Pecam1*, *Cd34*, *Tie1*, *Egfl7*). Trophoblast 2 cells expressed *Hsd11b2* (an enzyme produced by trophoblast cells (Togher et al., 2014)) and the prolactin receptor *Prlr*, which plays a role in trophoblast invasiveness and migration (Stefanoska et al., 2013). Trophoblast 3 cells expressed keratins (*Krt7*, *Krt8*, *Krt18, Prap1*, expressed by the uterine luminal epithelium (Diao et al., 2010)) and the early trophoblast marker *Wfdc2* (Yabe et al., 2016). Finally, VSMCs expressed actin-binding and contractile genes such as *Acta2*, *Myl9*, *Cnn1* and *Tagln* (**Fig. 7b**).

Cell subset composition was similar across different mice and genotypes (**Fig. S7e-g**), but we identified IUGR-associated expression changes in cell intrinsic expression in multiple cell types, either shared across cell types, or specific to a subset (Pseudobulk differential expression analysis, **Fig. 7c**). For example, *Anxa2* (a member of the Annexin family which plays a role in cell migration, invasion and angiogenesis (Lokman et al., 2011)) was downregulated in cells from IUGR mice compared to controls in different cell types (**Fig. 7c**), which we validated *in situ* (**Fig. S7h**). There were cell type-specific differentially expressed genes in many non-NK subsets. These included IUGR-up or downregulated stromal cell genes with roles in angiogenesis (*Rnf213, Slit3*) (Kobayashi et al., 2015; Ohkubo et al., 2015), regulation of fetal growth (*Igf1, Igfbp2, Igfbp3, Igfbp4)* (Nawathe et al., 2016), invasion and migration (*Usp25, Prlr, Aspn, Spon2*) (Ding et al., 2019; Lu et al., 2020; Satoyoshi et al., 2015; Stefanoska et al., 2013), and decidualisation and Wnt signalling (*Sfrp4, Rnf213, Wnt4*) (Carmon and Loose, 2008; Scholz et al., 2016). Some of these genes have also been implicated in the pathogenesis of pre-eclampsia (*Sfrp4, Wnt4*) (Wang et al., 2016; Zhang et al., 2013) and small-for-gestational-age pregnancies (*Dcn, Igf1, Igfbp2, Igfbp3, Igfbp4*) (Murthi et al., 2016; Nawathe et al., 2016) (**Fig. 7c and S7i**). Macrophages and dendritic cells showed changes in genes including *Osm*, involved in invasion under hypoxic conditions (Wie et al., 2018), and transcription factor *E2f1* that affects dendritic cell maturation (Fang et al., 2010) and is also implicated in pre-eclampsia (Kaitu’u-Lino et al., 2015). Macrophage cell subsets expressed genes were enriched for dendritic cell maturation, antigen presentation or inflammation genes (FDR = 0.007, 0.02, 0.0001, IPA) (**Table S2**). Trophoblasts and vSMCs showed changes in genes involved in remodelling of ECM (*P4ha1*) (Gilkes et al., 2013), invasion (*Klf5*) (Ma et al., 2017), regulation of angiogenesis or ECM remodelling (*Klf5, Cyr61, Col4a2, Bmper*) (Babic et al., 1998; Heinke et al., 2008; Kamphaus et al., 2000; Shindo et al., 2002) and stress response (*Sparc*) (Fenouille et al., 2011) (**Fig. 7c and Fig. S7j**). Some of these differentially expressed genes have been associated with pre-eclampsia or IUGR (*Sparc, Rgs2, Cyr61*) (Gellhaus et al., 2007; Perschbacher et al., 2020; Tossetta et al., 2019; Zhang et al., 2010). This highlights how the interaction between maternal KIR2DL1 and paternal HLA-C*05 has profound effects on several placental non-NK cell types and underlines a role for these genes and cellular functions in the etiology of IUGR.

### Maternal KIR2DL1 and paternal HLA-C*05 lead to change in cell-cell interaction networks in IUGR

Finally, we analysed our data to identify changes in putative cell-cell interactions in IUGR *vs*. control mice. We identified putative interactions between cell subsets (including within the same cell type) based on expressed receptor-ligand pairs (**Fig. 7a, Methods**), scored significant interactions with CellPhoneDB in WT x C*05 control and KIR x C*05 IUGR groups separately (**Fig. 7d,e**), and highlighted the interactions which were increased (red) or decreased (blue) between any pair of cell subsets in the IUGR network (**Fig. 7e**).

At baseline, the highest number of significant interactions occurred between stromal cells, fibroblasts, trophoblasts, and vSMCs, and both cNKs and trNKs had significant interactions with most other cell types (**Fig. 7d**). 38 of the 399 cell subset pairs with significant interactions in at least one condition had at least 20% more significant receptor-ligand interactions in IUGR than in WT (some as many as 50% more). Conversely, only nine of the cell type pairs had fewer significant receptor-ligand interactions in IUGR (**Fig. 7e**). Because more cell subset pairs had an increase rather than a decrease in the number of significant interactions in IUGR, we hypothesise that there is an overall increase in predicted intercellular communication at the maternal-fetal interface in IUGR.

We next focused on putative receptor-ligand interactions between NK cells and the other cell types that were detected only in IUGR or only in controls (selected examples in **Fig. 7f**). For example, the putative interaction between *Il15r* on NK cells and *Il15* on trophoblast 3 subtype was found to be significant in IUGR, but was not significant in the control group; IL-15 has been suggested to impact stromal cell decidualization and regulation of uNK cell differentiation and function (Dunn et al., 2002; Okada et al., 2000; Ye et al., 1996). Furthermore, there is a significant interaction between *Spn* on NK cells and *Icam1* on trophoblast subtype 1 in IUGR, but not WT cells. Indeed, overexpression of ICAM-1 by the villous trophoblast cells has been shown in placentitis and thought to play a role in the rupture of the trophoblast barrier (Juliano et al., 2006). In another example, there is a significant interaction between *TgfβR1* on NK cells and *Tfgβ2* on the stromal or trophoblast cell subtypes in IUGR, consistent with TGFβ’s control of trophoblast invasion, proliferation and migration (Davies et al., 2016). Similarly, there is a significant interaction between *Notch1* in NK cells and *Dlk1* in trophoblast cells or SMCs in the IUGR mating but not the control group; the Notch pathway is one of the key pathways regulating angiogenesis (Zhao and Lin, 2012). Overall, these results highlight potential roles for specific putative interactions between uNK cells and other cell types that could contribute to the development of or be the result of IUGR.

## Discussion

IUGR is one of the most common obstetric diseases and is associated with perinatal morbidity and mortality, as well as several long-term medical conditions such as heart disease, type 2 diabetes and stroke. While the consequences of IUGR are well understood, much is unknown about the underlying mechanisms that cause it and effective screening and treatment strategies are needed. One major challenge in studying pathogenic mechanisms of pregnancy complications is a lack of accessible tissue at relevant points early in gestation, especially when the disease manifestations are seen only later in pregnancy; therefore, a physiologically relevant model system for studying IGUR is necessary. Furthermore, even though there is evidence that HLA and KIR genes confer risk to IUGR, these genes are highly polymorphic and inherited in complex haplotypes in the human population (Bashirova et al., 2006; Lanier, 2005; Parham, 2005; Trowsdale, 2001; Vilches and Parham, 2002), making it challenging to dissect the role of individual genes.

Here, using a stepwise and multidisciplinary approach, we demonstrate a functional basis for how specific risk genes inherited from both parents can, in combination, lead to IUGR. Using a novel humanised transgenic model, we provide direct evidence for the involvement of the inhibitory KIR receptor, KIR2DL1, expressed on maternal uNK cells, and its interaction with fetal HLA-C*05 in manifestation of IUGR. Detailed characterisation of the model by imaging of the uteroplacental vasculature demonstrated that this genetic risk translated to changes in uterine spiral artery diameter and segment length and therefore, a higher resistance or decreased blood flow to the fetuses leading to growth restriction, implicating the KIR-HLA gene interaction in remodeling of the uterine vasculature.

IUGR and pre-eclampsia may have a shared disease pathogenesis, although mechanistic differences between these conditions have not been widely studied. In our model, we observe IUGR, evidenced by reduced average fetal weight, a fetal weight distribution skewed towards < 5^th^ and 10^th^ percentile, and placentation defects caused by altered uterine spiral arteries — but we did not see symptoms of pre-eclampsia (*e.g.* hypertension or proteinuria). It is possible that the absence of pre-eclampsia in our model is because pre-eclampsia is a disease of the human species and does not seem to occur in most other mammals (Robillard et al., 2002); however, it is also possible that manifestation of pre-eclampsia requires additional risk factors or mechanisms. Thus, our model might serve as baseline for investigating which additional factors could lead on from IUGR to pre-eclampsia.

Characterisation of cell types from the maternal-fetal interface by scRNA-seq revealed cellular heterogeneity and function as well as gene programs involved in placentation and the development of disease in pregnancy. In particular, we built a single cell atlas of sorted uNK cells from both healthy controls and mice with IUGR, at a time point in gestation when uNK cells are most prevalent and contribute to important biological pathways in pregnancy. We identified seven major uNK subtypes and 16 cell states. The uNK subtypes play diverse roles at the maternal-fetal interface, demonstrated by their functional heterogeneity—ranging from cellular migration, regulation of NF-kB activity, to those involved in cell-cell and cell-matrix interactions and metabolic pathways. Topic modeling revealed fluidity in cell states where cells from different uNK subtypes participated in the same gene program or a proportion of cells from within a specific subtype participated in a gene program. While the uNK cells have been shown to be important in pregnancy, our findings reveal the complex roles these cells play at a critical timepoint in pregnancy both in terms of the different biological programs and the uNK subtypes.

Both cNK and trNK subtypes are involved in biological processes altered or deficient in IUGR, including chemokine signaling, degradation of ECM components, and protection against oxidative damage. The involvement of uNK cells in these processes, which play an important role in placentation and vascular remodeling, demonstrated that signaling via the inhibitory receptor KIR2DL1 dampened the function or response of uNK cells, thus contributing to development of IUGR. This is concordant with previous observations that uNK cells surround spiral arteries and release angiogenic factors that play a role in placentation and vascular remodeling (Croy et al., 2000). Further work could involve studying the role of specific genes within the biological processes deficient in disease.

Profiling of unsorted cells from the mouse maternal-fetal interface at mid-gestation showed that despite the NK specific expression of KIR genes, there are substantial transcriptional changes in the IUGR model in several non-NK cell types from the maternal-fetal interface—impacting key functions such as stromal cell decidualization, Wnt signalling, angiogenesis, invasion and ECM remodeling. We hypothesize that this is a result of cellular crosstalk between uNK cells and the cells in the microenvironment, possibly though receptor-ligand interaction pairs present exclusively in the IUGR mice or control mice, between uNK cells and other cell types.

The gene expression changes in cells at the maternal-fetal interface in IUGR were moderate in magnitude but spanned many genes and cell subsets. This combined effect is likely to be further amplified due to the sensitivity to changes to placentation and the fetus at this early and critical stage of gestation. This could imply that healthy pregnancy represents a fine tuned and highly calibrated system and a disturbance in that balance leads to a multitude of downstream effects ultimately manifesting in disease.

We saw striking similarities in genes and pathways that were differentially expressed in IUGR with those that play a role in tumour progression and metastasis. For example, processes such as cellular invasion, migration, angiogenesis, cellular proliferation and differentiation are important in placentation, but are also involved in malignancy. Unlike the uncontrollable growth that occurs in cancer, these processes, however, are normally tightly controlled in development of the placenta (Januar et al., 2015; West et al., 2018). Understanding of the mechanisms that perturb these processes in pregnancy disorders might provide insights into tumour metastasis and vice versa, and also provide a starting point for developing new drugs to intercept IUGR at the earliest possible timepoint. The safety of such drugs will clearly need to be stringent as side effects could be more severe than IUGR because of the sensitivity of the fetus.

Our data provide essential functional data to support human genetic studies associating maternal KIR AA genotype with risk for pregnancy complications when the fetal group 2 HLA-C genes are paternally inherited (Hiby et al., 2010). Specifically, we did not observe significant IUGR in the C*05/KIR x WT mating combination, where the group 2 HLA-C gene in the fetus was maternally inherited and the mother and the fetus had the same level or number of copies of group 2 HLA-C genes.

IUGR is difficult to predict and it has been found that growth restricted fetuses > 10^th^ percentile in size can remain undiagnosed despite not meeting their growth potential and having an increased risk for adverse perinatal outcome (Gordijn et al., 2016; Malhotra et al., 2019; Nardozza et al., 2017). Therefore, identifying risk genes and early prenatal events that lead to IUGR are crucial. Identifying specific KIR and HLA-C as risk genes and an understanding of how they mediate disease means that they could be used to screen parents and identify infants with increased risk of neonatal complications, thereby having the potential to allow development of new treatment modalities and improve pregnancy management and clinical outcomes for both the mother and the baby.

## Acknowledgments

L.F. was supported by the Wellcome Trust (grant no. 100308/Z/12/Z), Danish National Research Foundation, Takeda, the Medical Research Council (grant no. MC_UU_12010/3), the Oak Foundation (grant no. OCAY-15-520) and the NIHR Oxford BRC. G.M. was supported by the Wellcome Trust (grant no. 100956/Z/13/Z) and the Li Ka Shing Foundation. We would like to thank V.Sexl (University of Veterinary Medicine of Vienna) for the NCR1-iCre transgenic mice, P.Höglund (Karolinska Institutet) for H2-Kb H2-Db knockout mice, B.Davies (Wellcome Trust Centre for Human Genetics, University of Oxford) for transgenic mouse generation services, D.Bowman, A.Ortiz and S.Jerman (Indica Labs, Albuquerque, NM, USA) for RNAscope image analysis services, L. Gaffney for figure edits, G.Holländer and M.Deadman (Dept. of Paediatrics, University of Oxford) for help with TEC isolations, C.Smillie, S. Simmons, A.Haber (Broad Institute of MIT and Harvard) and A.-C.Villani (Harvard Medical School) for brainstorming and discussions, G.Douglas and V.Rashbrook (Radcliffe Dept. of Medicine, University of Oxford) for help with blood pressure measurement in mice, A. Vernet (University of Oxford) for helping with micro-CT scanning, R.Kuehn (Helmholtz Center Munich) for help with transgenic vector design. We would like to acknowledge S-A.Clark, C.Waugh, K.Clark and P.Sopp in the flow cytometry facility at the MRC WIMM for providing cell sorting services. The flow cytometry facility is supported by the MRC HIU; MRC MHU (MC_UU_12009); NIHR Oxford BRC; Kay Kendall Leukaemia Fund (KKL1057), John Fell Fund (131/030 and 101/517), the EPA fund (CF182 and CF170) and by the MRC WIMM Strategic Alliance awards G0902418 and MC_UU_12025. We also thank the MRC WIMM core Transgenic service team for providing cryopreservation services. This project has been funded in part with federal funds from the Frederick National Laboratory for Cancer Research, under Contract No. HHSN261200800001E. The content of this publication does not necessarily reflect the views or policies of the Department of Health and Human Services, nor does mention of trade names, commercial products, or organizations imply endorsement by the U.S. Government. This Research was supported in part by the Intramural Research Program of the NIH, Frederick National Lab, Center for Cancer Research.

## Author Contributions

Conceptualization, G.K. and L.F.; Methodology, G.K., C.B.M.P., O.A., S.J.R., M.H., A.S., I.A.D. and G.M.; Software, C.B.M.P., G.K., J.L., O.A., S.J.R., M.H. and A.S.; Formal Analysis, C.B.M.P., G.K., M.A., J.L., O.A., S.J.R., M.H., A.S., K.E.A., C.D. and J.D.; Investigation, G.K., C.B.M.P., M.A., S.B.K., K.E.A., C.D., H.G.E., L.T.N., D.D., A.N., L.T.J., T.B. and E.S.; Writing – Original Draft, G.K. and C.B.M.P.; Writing – Review & Editing, G.K., C.B.M.P., O.A., A.R. and L.F.; Funding Acquisition, L.F., A.R., O.R.-R. and G.K.; Supervision, L.F., A.R., O.R.-R., M.C.; Project Administration, G.K., C.B.M.P., O.R.-R. and L.F.

## Supplementary Figure Legends

**Fig. S1.**
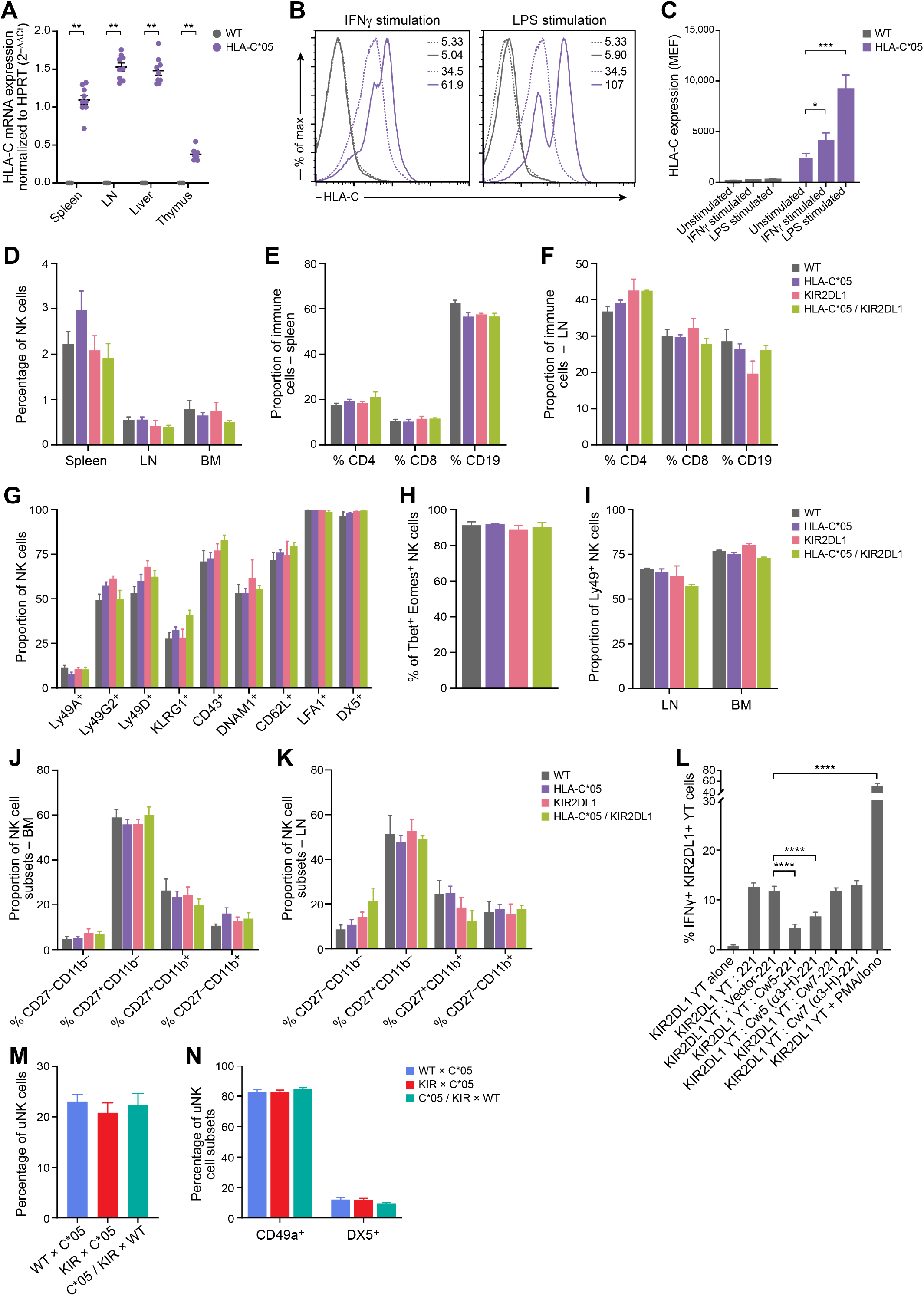
Functional characterisation of the HLA-C*05 and KIR2DL1 transgenic mouse model. (**a**) HLA-C*05 mRNA expression using tissue from different organs from WT or HLA-C*05 transgenic mice. Data is shown normalised to a HLA-C*05 spleen sample. (**b**) Representative HLA-C*05 cell surface expression on total spleen cells cultured with or without stimulation. Dashed lines represent unstimulated cells and solid lines represent stimulated cells. HLA-C*05 and WT mice are shown in purple and grey respectively. Numbers denote MFI. (**c**) HLA-C cell surface expression plotted as MEF of total splenocytes cultured with or without stimulation. (**d**) Percentage of NK cells (CD3^−^TCR^−^ NKp46^+^ cells) in spleen, LN and bone marrow (BM) from the different transgenic mice. (**e and f**) Percentage of total CD3^+^CD4^+^, CD3^+^CD8^+^ and CD19^+^ cells in the spleen and LN in the different transgenic mice. (**g**) Percentage of splenic NK cells expressing the different NK cell surface receptors/maturation markers as highlighted in the plot (percentages of receptor expression is gated on CD3^−^TCR^−^NKp46^+^ NK cells). (**h**) Percentage of splenic NK cells expressing transcription factors T-bet and Eomes in the different transgenic mice. (**i**) Proportion of Ly49^+^ NK cells (stained using antibody clones 14B11 and 4D11) in the LN and BM in the different transgenic mice. (**j and k**) Cell surface staining of CD27 and CD11b on gated NK cells from the bone marrow (**j**) or LN (**k**) from the different transgenic mice. (**l**) Response of KIR2DL1-expressing YT NK cell line (measured by IFN-*γ* production by flow cytometry) when co-cultured with 721.221 cells alone or 721.221 cells transduced with lentiviral HLA-C*05, HLA-C*07 or vector-transduced cells or with stimulation with PMA and Calcium Ionomycin. Additional controls included co-culture with 721.221 cells that were transduced with HLA-C*05 and HLA-C*07 constructs where the HLA *α*3 domain, transmembrane and cytoplasmic regions were cloned in from the respective HLA-C alleles (see **Methods** for details). (**m**) Percentage of uterine NK cells (CD3^−^CD19^−^TCR^−^CD45^+^NKp46^+^CD122^+^) at the maternal-fetal interface at gd9.5 are shown from the different transgenic crosses. Mating crosses are mentioned as female x male. (**n**) Percentage of CD49a^+^ or DX5^+^ uNK cell subsets at gd9.5 from the different transgenic crosses. Crosses are mentioned as female x male. Mean ± SEM is shown. (**a**) n= 3-9 per group. (**c**) n = 4-7 per group. (**d**) n = 3-11 per group. (**e**) n = 4-16 per group. (**f**) n = 3-10 per group. (**g**) n = 3-12 per group. (**h**) n = 3-10 per group. (**i**) n = 3-9 per group. (**j and k**) n = 3-10 per group. (**l**) n = 8-9 per condition (**m and n**) n = 8-10 per group. *p < 0.05, **p < 0.01, ***p < 0.001, ****p < 0.0001, Mann-Whitney test.

**Fig. S2.**
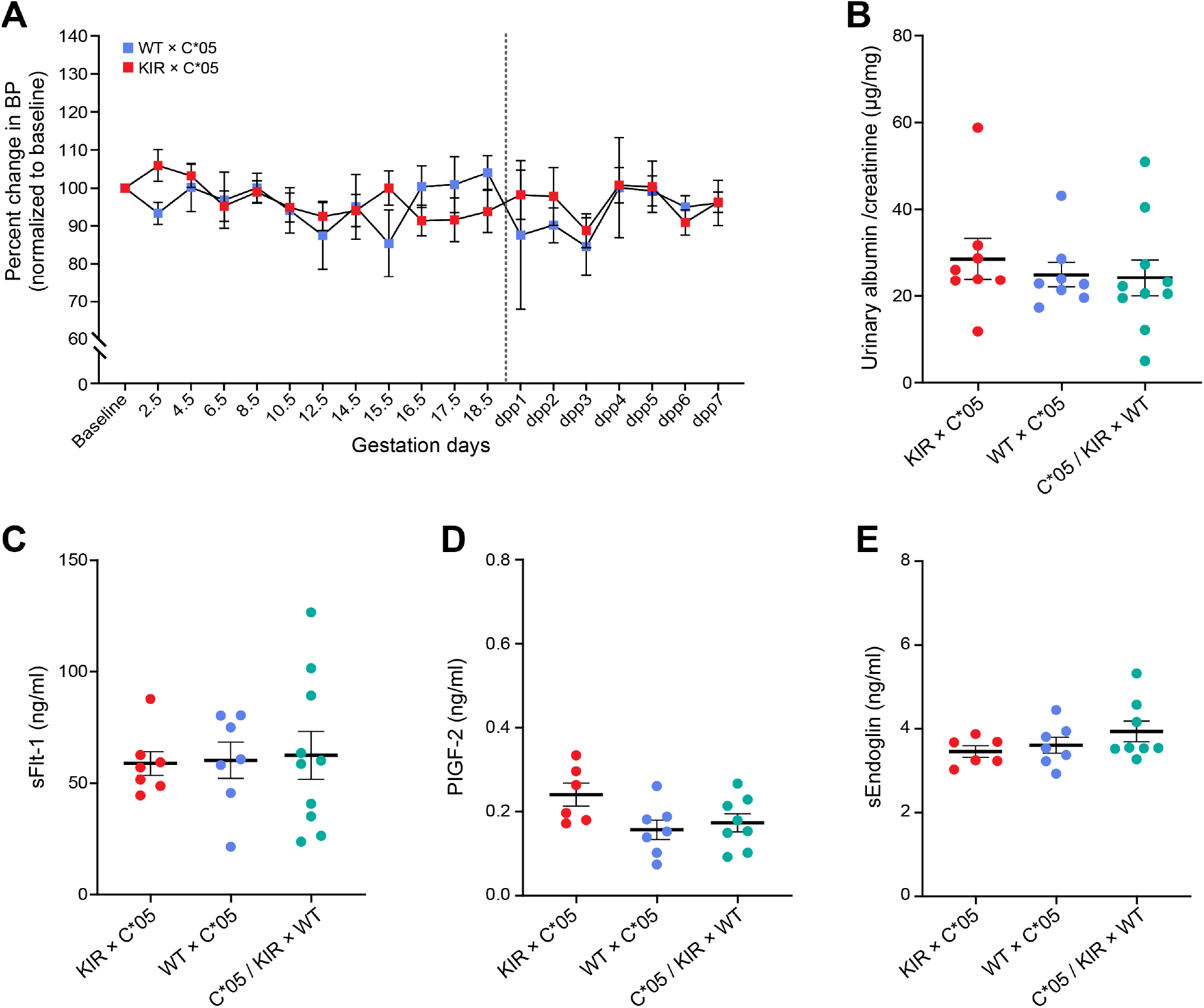
Assessment of blood pressure, urinary protein and plasma proteins. (**a**) Systolic blood pressure measured using non-invasive tail-cuff photoplethysmography during pregnancy and post-partum in the depicted transgenic mating crosses. Grey line indicates birth of the pups. Blood pressure at each time point is shown relative to the baseline for each mouse (measured pre-pregnancy). (**b**) Assessment of protein in urine samples collected from gd18.5 pregnant mice from the different transgenic mating crosses. Urinary albumin/creatinine is shown. (**c, d and e**) Plasma concentrations of sFlt-1 (**c**), PIGF-2 (**d**), sEndoglin (**e**) in pregnant gd18.5 mice from the different transgenic mating crosses. (**a**) n = 7-9 per group. Not all mice gave valid blood pressure measurements on every measurement day. Represented data includes observations from a minimum of 3 mice per group on any measured day. (**b**) n = 8-10 per group. (**c**) n = 7-10 per group. (**d and e**) n = 6-8 per group. Mean ± SEM is shown. Mann-Whitney test.

**Fig. S3.**
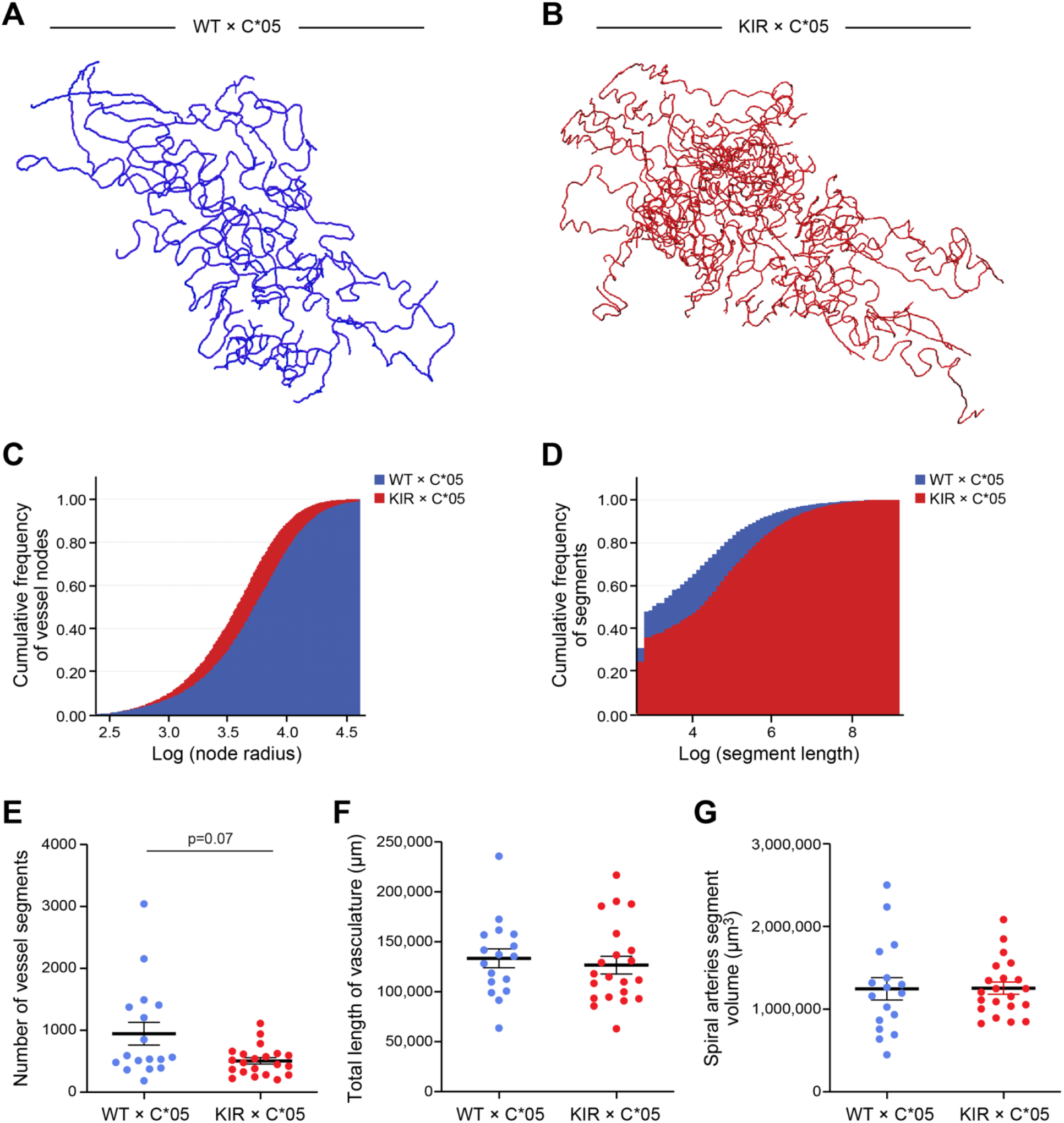
Changes in uterine spiral arteries during gestation. (**a and b**) Representative skeletonised vascular network from the segmentation analysis from the WT x C*05 and KIR x C*05 mating crosses at gd10.5 are shown. (**c and d**) Cumulative distributions of vessel radii nodes (**c**) and segment lengths (**d**) from the mating crosses are depicted. (**e**) The number of spiral artery vessel segments are shown. (**f and g**) Total length of vasculature (**f**) and volume (**g**) measured from the spiral artery segmented data from WT x C*05 and KIR x C*05 mating crosses is shown. Data is represented as Mean ± SEM. (**c - g**) n = 17–21 implantation sites per group. Mann-Whitney test.

**Fig. S4.**
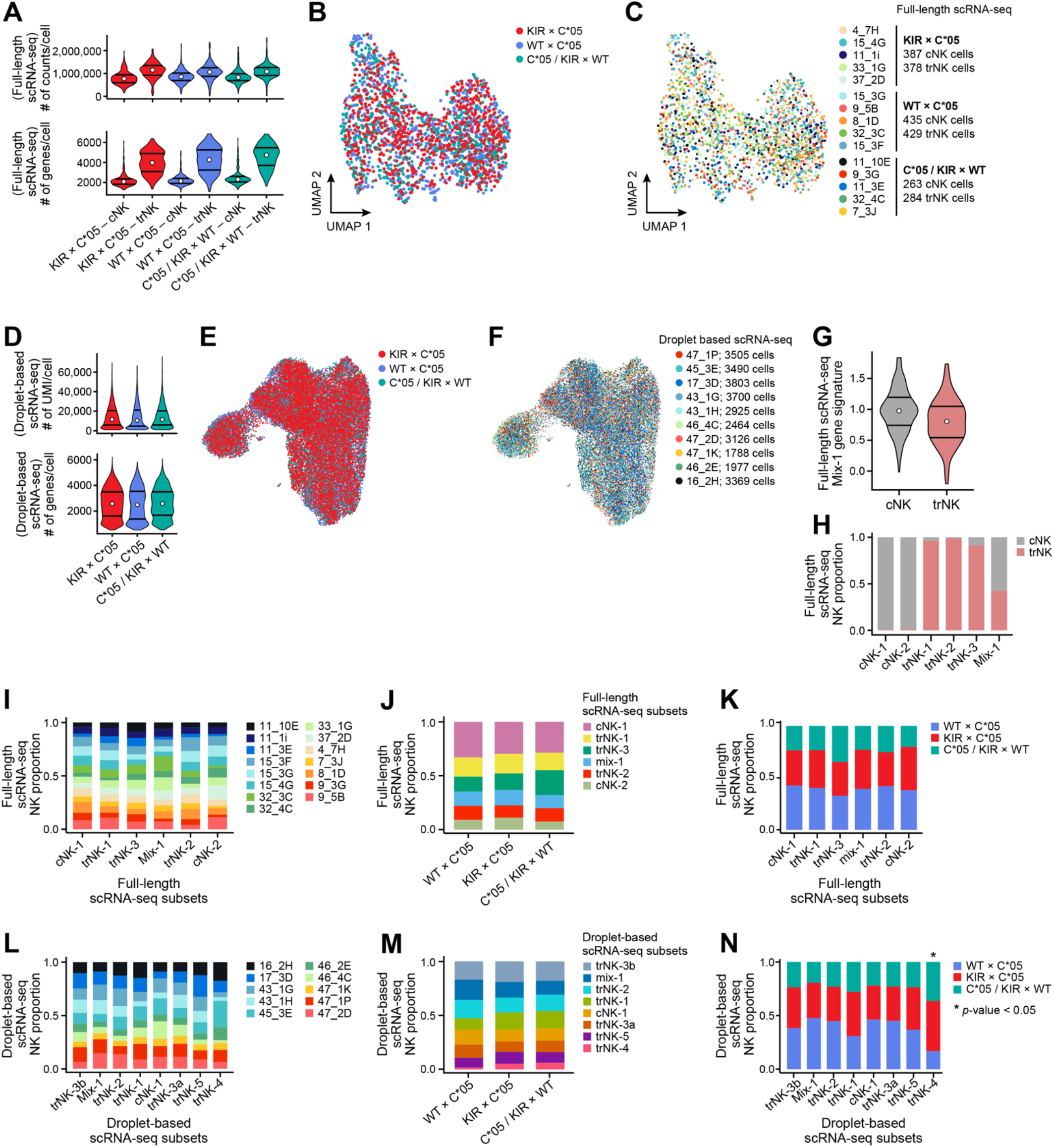
Quality control metrics for full length and droplet-based scRNA-seq data of sorted NK cells at the mouse maternal-fetal interface. **(a)** Distribution of the number of counts per cell and the number of genes per cell in each genotype for the full length sorted NK data. The first and third quartiles and median are indicated on each violin plot. The genotypes are displayed along the x axis, and conventional NK cells are indicated by “_cNK” and tissue resident NK cells are indicated by “_trNK”. **(b-c)** UMAP embedding of sorted NK cells from the maternal fetal interface, analyzed by full length scRNA-Seq. Colors of the dots correspond to genotype (**b**) or the mouse of origin (**c**). **(d)** Distribution of the number of UMI per cell, the number of genes per cell in each genotype from the droplet based scRNA-seq uNK data. The first and third quartiles and median are indicated on each violin plot. **(e-f)** UMAP embedding of sorted NK cells analyzed by droplet based scRNA-seq, with cells colored by genotype (**e**) or mouse of origin (**f**). (**g**) Cells expressing the gene signature from the mix-1 cell subset from the full length scRNA-seq data stratified into cNK or trNK cells based on FACS sorting. (**h**) Proportion of cNK and trNK cells (defined by FACS) in each cell subset for the full length scRNA-seq NK dataset. (**i-k**) Proportion of composition of clusters depicted in Fig. 5B as per individual mice (**i**) or genotype (**j, k**). (**l-n**) Proportion of composition of clusters depicted in Fig. 5D as per individual mice (**l**) or genotype (**m,n**). p-values represent whether the number of cells belonging to the IUGR phenotype (KIR x C*05 mating) are significantly more or less than the two control phenotypes, and were calculated using a latent Dirichlet allocation model.

**Fig. S5.**
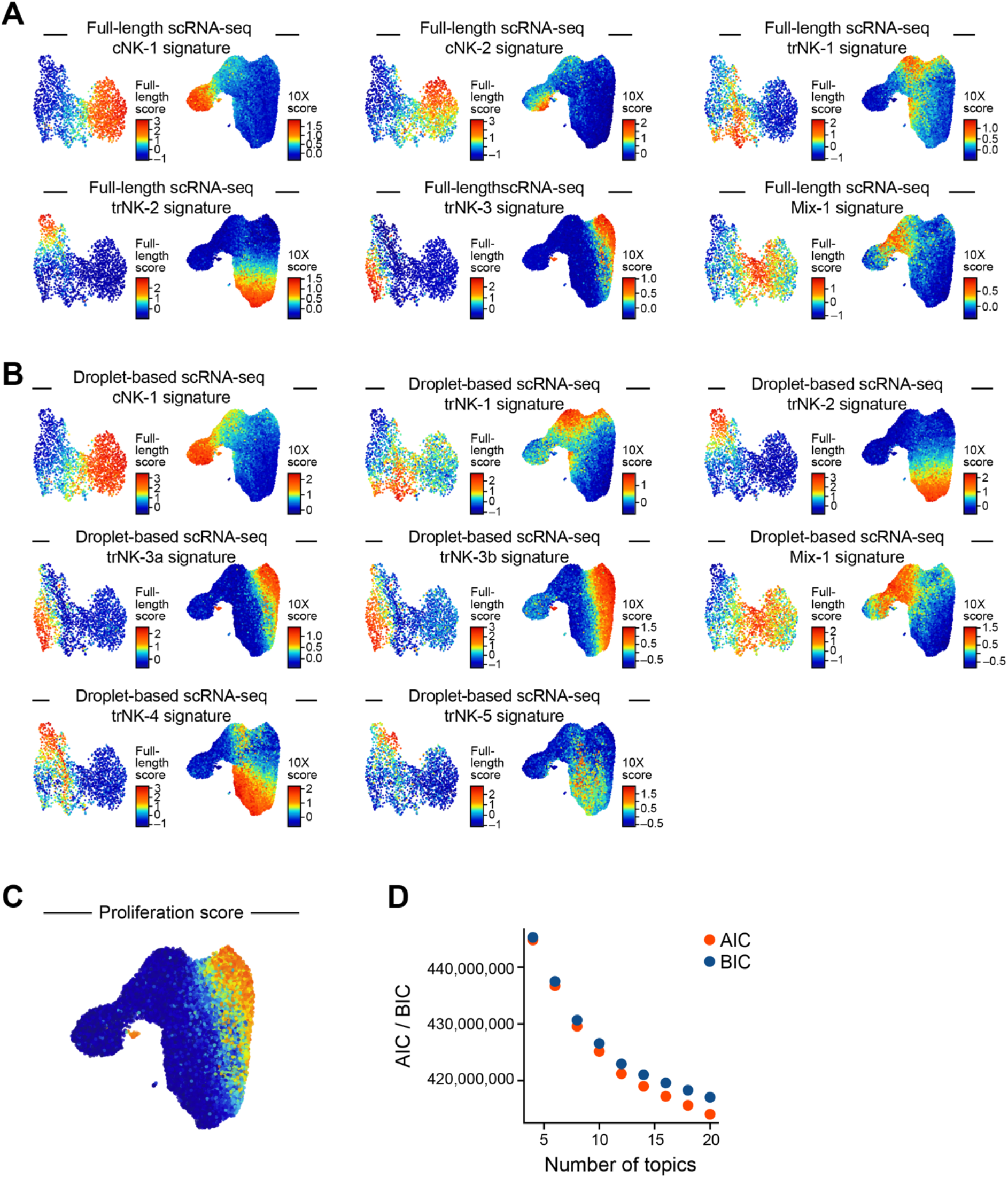
Comparison of NK subtype signatures observed between the full length and droplet based scRNA-seq datasets. **(a)** NK subtype (cluster) signatures from full length scRNA-Seq data, plotted on UMAP embeddings for both full length and droplet based scRNA-seq data sets to highlight consistency in observed heterogeneity between the two data sets. Red indicates a cell has a high signature score and blue indicates the cell has a low signature score. **(b)** NK subtype (cluster) signatures from the droplet based scRNA-seq data set, plotted on UMAP embeddings for both full length and droplet based scRNA-seq data sets (the reverse of **b**). **(c)** UMAP embedding of droplet based scRNA-seq NK data, colored by proliferation score. **(d)** Akaike information criterion (AIC) and Bayesian information criterion (BIC) for iterations of topic modeling where k was set to 4:20 in increments of 2; this information was used to help select the value of k to use for our final topic modeling analysis.

**Fig. S6.**
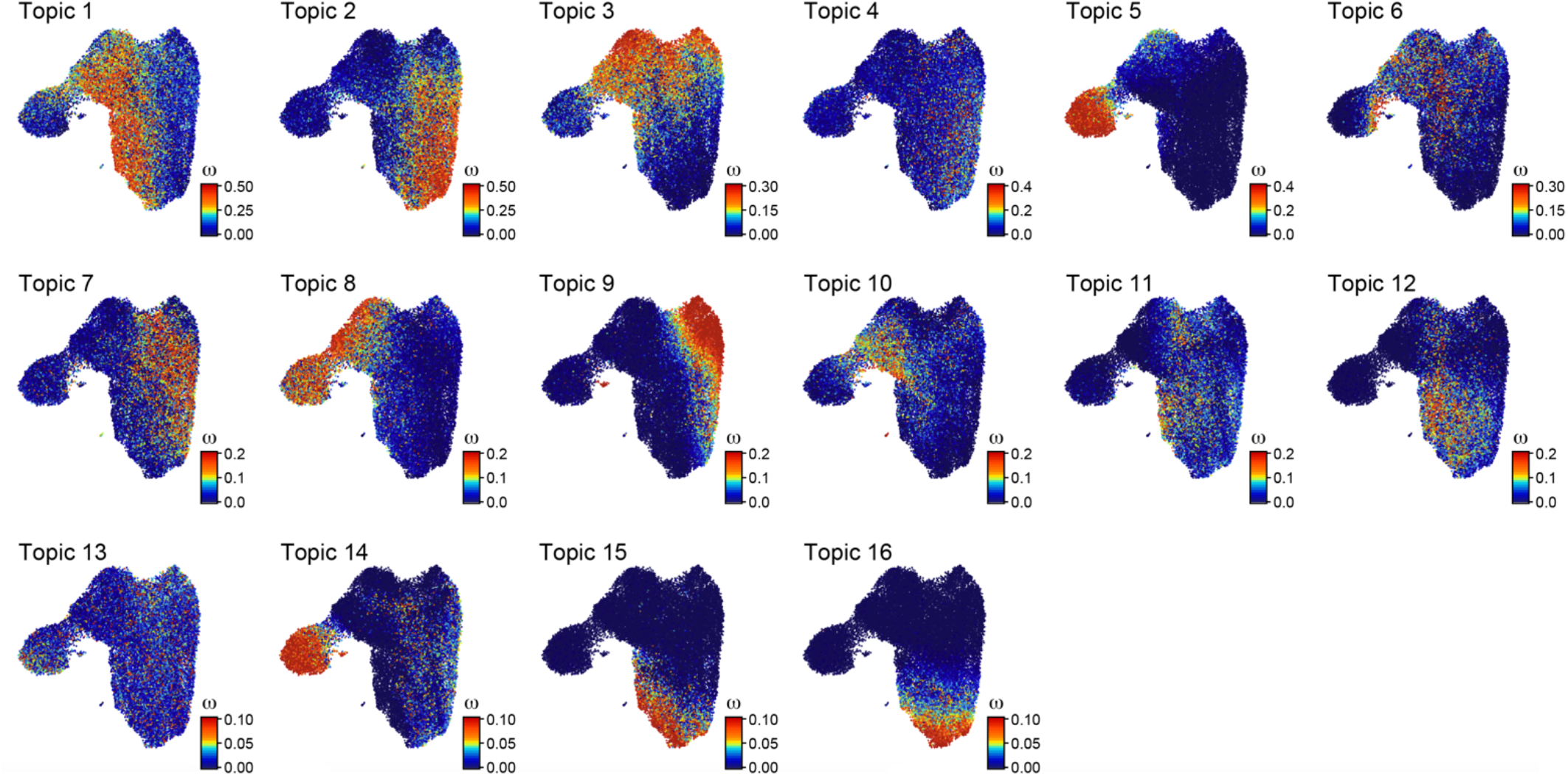
Topic modeling of droplet based scRNA-seq NK cell data from the maternal-fetal interface. Topics from topic modeling analysis depicting all 16 topics. UMAP embedding colored by the cell weight for that topic – bright red indicates the cell is high in the topic and dark blue indicates the cell is low in the topic.

**Fig. S7.**
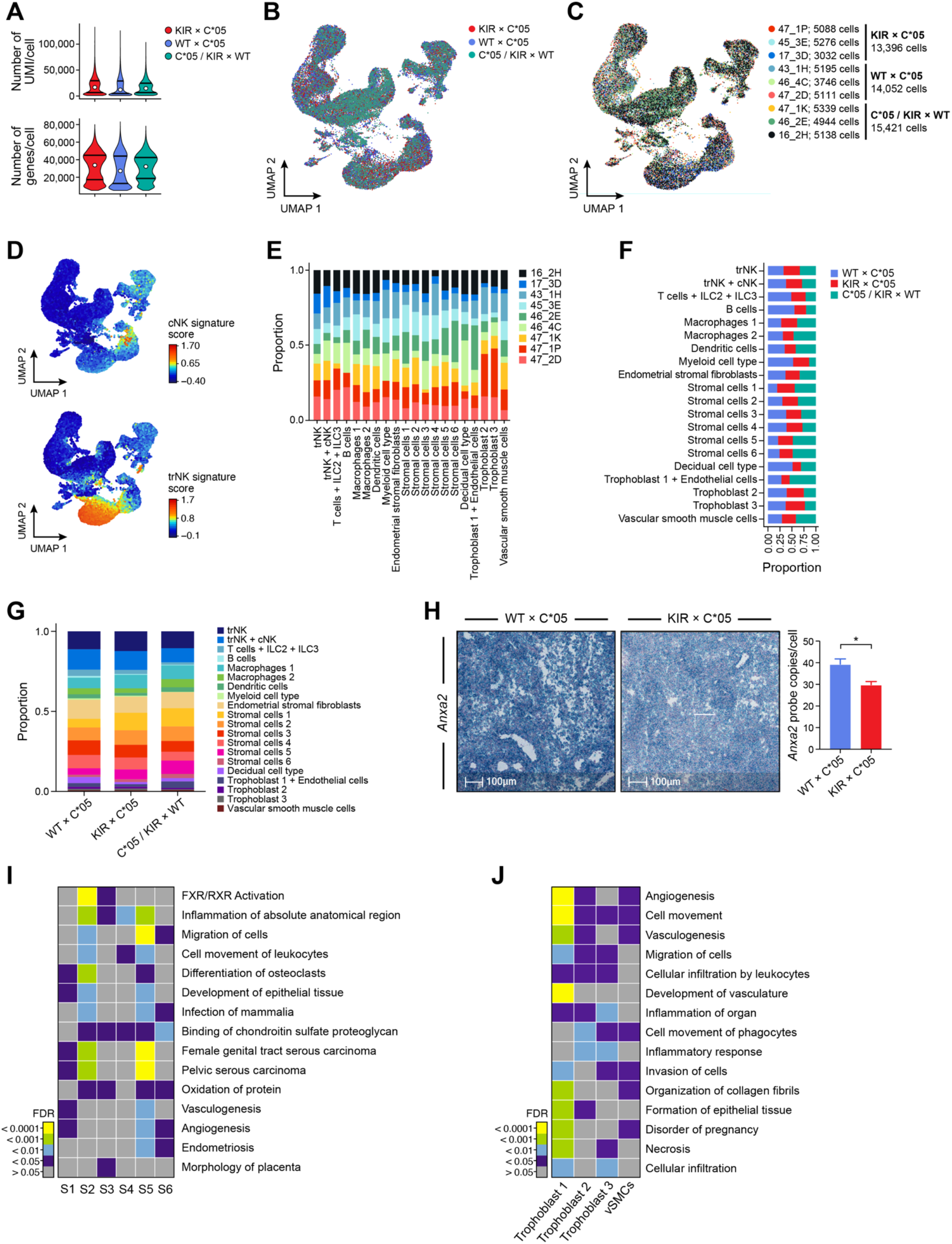
Quality control metrics for droplet based scRNA-seq data of unsorted cells at the mouse maternal-fetal interface and differentially expressed pathways. **(a)** Distribution of the number of UMI per cell and the number of genes per cell in each genotype, for droplet based scRNA-seq data from unsorted cells. **(b-c)** UMAP embedding of unsorted cells analysed by droplet based scRNA-seq colored by genotype (**b**) or mouse of origin (**c**). **(d)** UMAP embedding of droplet based scRNA-seq data from unsorted cells colored by cNK and trNK signature score. (**e**) Proportion of composition of clusters/cell types depicted in Fig. 7b stratified by individual mice (**e**) or genotype (**f,g**). **(h)** Re-validation of *Anxa2* using *in-situ* hybridisation (RNAscope) on uterine tissue collected from implantation sites at gd9.5. Representative probe staining on sections from WT x C*05 or KIR x C*05 implantation sites is shown. *Anxa2* staining is seen in teal. Quantification of probe staining depicted as average probe copies per cell within a stained section is shown. n=10 sections, Mann-Whitney test. (**i,j**) Comparison of pathways and functions altered in stromal cell types (**i**), Trophoblast and SMCs (**j**) in KIR x C*05 IUGR mice compared to the controls, assessed using Ingenuity pathway analysis. Colors of heatmap indicate BH-FDR value of enrichment.

## Supplementary Tables

**Table S1 – Signatures used in QC**

**Table S2 – Pathway analysis using IPA on differentially expressed genes on selected cell type clusters from droplet-based unsorted cells data**

**Table S3 – List of oligonucleotides**

## METHODS

### Animals

All animal experiments were approved by the local Ethical Review Committee at the University of Oxford, and performed under license from the UK home office (project license numbers 30/3386 and P0A53015F) in accordance with the Animals (Scientific Procedures) Act, 1986. Mice were maintained in a pathogen-free facility in individually ventilated cages in a temperature- and humidity-controlled room with a 12h light/12h dark cycle under standard housing conditions with continuous access to food and water. Male and female mice were used across different experiments. None of the mice had any noticeable health abnormalities. For timed-mating experiments, male and female mice were mated in the afternoon of one day and checked for plugs and separated on the morning of the next day. Detection of the plug was considered as gd0.5. In addition, female mice were weighed prior to being set-up in timed-mating experiments, and on every alternate day from gd7.5 onwards in order to confirm pregnancy. These refinements avoided wastage of any pregnant transgenic mice.

### Generation of transgenic mice

The HLA-C*05 and KIR2DL1 transgenic mice were generated using targeted insertion into the *ROSA26* locus. For generation of the HLA-C*05 transgenic mice, to facilitate interaction of HLA-C with murine CD8 (and hence the T cell receptor), the *α*3 domain of HLA-C*05 was replaced with its murine counterpart from H2-K^b^, along with the adjacent transmembrane and cytoplasmic domains. To allow for this, the *HLA-C*05* construct was made by amplifying ∼2.04 kb genomic fragment of *HLA-C*05:01:01:01*, which contained 776 bp of the *HLA-C* 5’UTR and exons 1-3 up to a midpoint in intron 3, which was fused to a ∼3.58 kb fragment of the genomic *H-2K^b^* gene, beginning at a midpoint in intron 3 and containing exons 4-8 and the *H-2K^b^* 3’UTR. The genomic DNA construct was first assembled in the pTZ18U vector (Sigma-Aldrich) prior to subcloning into vector pCB92 (Chen et al., 2011), downstream of a promoterless neomycin selection cassette and a murine H19 insulator, creating plasmid CB92-C*05 in which the whole transgene array is flanked by PhiC31 attB sites.

For generation of the KIR2DL1 transgenic mice, the coding DNA sequence of *KIR2DL1*0030201* along with a Kozak consensus sequence (GCCACC) immediately upstream of the ATG start codon, was amplified and cloned into the pEx-CAG-stop-bpA vector (a gift from Ralf Kühn), between the loxP flanked STOP cassette and the polyadenylation (pA) signal. The presence of the STOP cassette (a puromycin resistance coding region followed by triple pA signals) served as the transcriptional STOP signal for transgene expression. The KIR2DL1 protein is expressed from the CAG promoter upon the Cre mediated excision of the loxP flanked stop cassette. This vector also included the promoterless neomycin selection cassette and PhiC31 attB sites required for PhiC31 integrase mediated cassette exchange.

For targeted insertion at the *Gt(ROSA26)Sor* locus, a PhiC31 integrase mediated cassette exchange approach was adopted using IDG26.10-3 ES cells, which are a (C57BL/6J x 129S6/SvEvTac) F1 ES cell line harbouring a PGK promoter driven hygromycin selection cassette flanked by PhiC31 attP sites, positioned within intron 1 of *Gt(ROSA26)Sor* (Hitz et al., 2007). The targeted insertion of the vector into the *ROSA26* locus ensures reproducible transgene expression. 1×10^6^ IDG26.10-3 ES cells (Hitz et al., 2007) were co-electroporated with 5 µg of either pCB92-C*05 or pEx-CAG-KIR2DL1 and 5 µg of pPhiC31o (Addgene) using the Neon transfection system (Thermo Fisher) (3×1400 V, 10 ms) and plated on G418 resistant fibroblast feeder layers. After approximately 7 days of selection in 350 µg/ml G418, 16 resistant colonies were isolated per construct, expanded and screened for the correct cassette exchange event at the 5′ and 3′ ends using specific screening primers provided in Table S3 (5′ screen: yields a 280 bp product; 3′ screen: yields a 518 bp product). Correctly integrated ES cell clones were injected into mouse C57BL/6J blastocysts, and the resulting chimeric males were mated to C57BL/6J females and the progeny were screened for germline transgene transmission, normal breeding and appropriate expression of the transgene. The mice were further backcrossed onto C57BL/6J for 4-5 generations. Mice at the same backcrossing generation were used as transgene-negative littermates or controls within each experiment. The NK1.1 antigen is not expressed on the 129 background. The NK1.1 antigen status was always assessed when performing matings and the NK1.1 expression was kept consistent within experiments with mice largely being maintained on a NK1.1-ve background. KIR2DL1-floxed transgenic mice were mated to NCR1-iCre mice (a gift from Veronika Sexl) (Eckelhart et al., 2011) to obtain KIR2DL1-NCR1-iCre transgenic mice which had specific expression of KIR2DL1 in NCR1-expressing cells, and referred to as KIR2DL1-expressing mice hereafter. KIR2DL1-NCR1-iCre transgenic mice were crossed to HLA-C*05 transgenic mice to generate double transgenic mice having expression of both HLA-C*05 and KIR2DL1. For adoptive transfer experiments, HLA-C*05 mice were mated to H2-K^b^H2-D^b^ double-knockout mice (a gift of Petter Höglund) to generate HLA-C*05 mice which lacked expression of H2-K^b^ and H2-D^b^. Genotyping of all alleles was done using PCR as described below.

### Cell lines

The B-lymphoblastoid cell line, 721.221 was purchased from the International Histocompatibility Working Group and grown in RPMI-1640 supplemented with 15% heat-inactivated fetal calf serum (FCS) (Sigma-Aldrich), sodium pyruvate (Thermo Fisher), L-glutamine (Sigma-Aldrich) and primocin (Invivogen). YT cells transfected with KIR2DL1 (a gift of Dr. Chiwen Chang) were grown in RPMI-1640 supplemented with 10% heat-inactivated FCS, L-glutamine and penicillin-streptomycin (Sigma-Aldrich). HEK 293T cells (ATCC) were cultured in DMEM supplemented with 10% heat-inactivated FCS, L-glutamine and penicillin-streptomycin.

### Genotyping of mice

Genomic DNA from mice and embryos was extracted using E-Z 96^®^ Tissue DNA Kit (Omega Bio-tek) as per the manufacturer’s instructions. The expression level of genes of interest for each sample was normalized against mouse housekeeping gene, Glyceraldehyde 3-phosphate Dehydrogenase (GAPDH). Genotyping of HLA-C*05, KIR2DL1, NCR1 and NK1.1 was performed using real-time quantitative PCR (qPCR) on a LightCycler 480 II (Roche) under the following conditions: 50°C for 2 min, 95°C for 10 min, 40 × Quantification mode (95°C for 15 s, 60°C for 1 min), Melting Curve mode (95°C for 15 s, 60°C for 15 s, 95°C Continuous Acquisition of 5 per °C), 40°C for 10 min. qPCR samples were prepared with Power SYBR Green Master Mix (Thermo Fisher) and gene-specific primers as listed in Table S3. Comparative CT method (2^−ΔΔCT^) was used for the quantitative analysis of relative gene expression, where ΔCT is the difference between the threshold cycle of gene of interest and GAPDH, and ΔΔCT is the difference between the ΔCt of each sample and the positive control sample. Genotyping of H2-K^b^ H2-D^b^ double-knockout mice was performed using Touchdown (TD) PCR on a SimpliAmp Thermal Cycler (Thermo Fisher) under the following conditions: 95°C for 10 min, 17 cycles of (95°C for 30 s, 60°C decreased by 0.5°C every cycle for 40 s, 72°C for 30s), 16 cycles of (95°C for 30 s, 60°C for 40 s, 72°C for 30s), 72°C for 10 min. TD-PCR samples were prepared with AmpliTaq Gold™ DNA Polymerase, Gold Buffer, MgCl2, GeneAmp™ dNTP Blend (Thermo Fisher) and gene-specific primers as listed in Table S3. TD-PCR products were electrophoresed in 1.2% Tris-Borate-EDTA (TBE) agarose gels, stained with Midori Green Advance DNA Stain (Geneflow) and imaged with Gel Doc XR+ Gel Documentation System (Bio-Rad).

### Preparation of single cell suspensions from mouse organs for phenotypic and functional characterisation experiments

All organs were placed in R10 medium (details below) on ice until further processing. Single cell suspensions were counted using an improved Neubauer chamber or Countess I (ThermoFisher) after dilution in 0.4% trypan blue (ThermoFisher). All centrifugation steps were done at 480g for 5 min at 4°C unless otherwise stated. R10 constituted of RPMI-1640 media supplemented with L-glutamine, sodium bicarbonate (Sigma-Aldrich), 10% heat-inactivated FCS, 1x Penicillin-Streptomycin and 50μM 2-Mercaptoethanol (Gibco), freshly added. Spleens were carefully dissociated using the plunger of a 5ml syringe and filtered through pre-wetted 70μm cell strainers (Corning). Cells were washed once in R10 and red blood cells lysed in Red Blood Cell Lysing Buffer Hybri-Max™ (Sigma-Aldrich) for 7 min at room temperature. Cells were washed in R10 and counted. Splenocytes were either used for cell surface and intranuclear flow cytometric staining or processed for *in vitro* stimulations. In some experiments, cells were also resuspended in RLT buffer (Qiagen) supplemented with 2-Mercaptoethanol and stored at −80°C until RNA isolation. For isolation of cells from lymph nodes, left and right inguinal and axillary lymph nodes were carefully dissociated using the plunger of a 5ml syringe and filtered through pre-wetted 70μm cell strainers. Cells were washed in R10, counted and then stained as per the flow cytometry protocol. Bone marrow isolations were done by collecting left and right tibia and flushing bones with 5ml Hanks Balanced Salt solution (HBSS) (Sigma-Aldrich) using a syringe with a 26G needle. Cells were dissociated by passing the bone marrow through the needle 5-10 times and filtered through pre-wetted 70μm cell strainers. Cells were washed once in R10 and red blood cells lysed in Red Blood Cell Lysing Buffer Hybri-Max™ for 7 min at RT. Cells were washed in R10, counted and then stained as per the flow cytometry staining protocol. Isolation of thymocytes were performed by dissociating thymi using the plunger of a 5ml syringe and filtered through pre-wetted 70μm cell strainers. Cells were washed in PBS, counted and then stained as per the flow cytometry staining protocol.

### Flow cytometric staining

Surface staining was done in the presence of anti-CD16/32 antibodies to block FcγRII/III receptors using Mouse Fc block^TM^ (BD Biosciences). Cells were incubated with the Mouse Fc block^TM^ for 10 min followed by incubation with fluorochrome-conjugated pre-titrated antibodies for 20 min at 4°C. Cells were washed in FACS buffer (PBS with 2% Heat-inactivated FCS) and fixed in 1x BD cellFIX (BD Biosciences). Samples were acquired on the LSRFortessa (BD Biosciences), Fortessa X-20 (BD Biosciences), Cyan ADP (Dako) or the Attune acoustic focusing cytometer (Applied Biosystems), and analysed using the FlowJo software (FlowJo). In order to ensure comparability, FluoroSpheres (Dako) and Sphero^TM^ Rainbow Calibration beads (BD Biosciences) both with defined MEF (molecules of equivalent fluorochromes) were run for each experiment in addition to the samples and used to calculate normalised MFI values. For intracellular staining, cells were fixed, permeabilised and stained using the BD Cytofix/Cytoperm kit (BD biosciences) as per manufacturer’s instructions. For nuclear staining of transcription factors, cells were stained using the Foxp3 staining buffer set (eBioscience) according to the manufacturer’s instructions. Samples were acquired and analysed as described above.

### Stimulation of splenocytes

For *in vitro* stimulation of total splenocytes, 2×10^6^ splenocytes were stimulated with 4μg/ml LPS (Sigma-Aldrich) or 50U/ml IFNγ (Peprotech) for 24 hours in 24-well plates in R10 medium. After incubation, cells were detached by careful pipetting, washed in FACS buffer and stained as per the flow cytometric staining protocol. For *in vitro* stimulation of NK cells, 1×10^6^ splenocytes were isolated from mice and cultured on Immulon U-bottom plates coated with antibodies for Ly49D (0.5mg/ml) (Biolegend) or NKp46 (1mg/ml) (Thermo Fisher) with or without the presence of anti-KIR antibody (1mg/ml) (Biolegend), in the presence of 2000 U/ml rhIL-2 (Peprotech) and Brefeldin A (10μg/ml) (Sigma-Aldrich) for 5 hours. Cells were then harvested and stained as per the flow cytometric staining protocol and IFN*γ* production by NK cells was assessed. Negative and positive controls included cells cultured without any stimulus and those cultured with PMA (250ng/ml) (Sigma-Aldrich) and Calcium Ionomycin (2.5μg/ml) (Sigma-Aldrich).

### Adoptive transfer experiments

Adoptive transfer experiments to assess *in vivo* rejection were performed as previously described (Oberg et al., 2004). Splenocytes were isolated from mice lacking expression of murine MHC class I i.e. H2K^b-^H2D^b-^ (referred to as KO cells), WT mice (which were H2K^b+^H2D^b+^) or HLA-C*05 expressing mice which had been bred to mice lacking expression of murine MHC class I molecules (i.e. HLA-C*05^+^ H2K^b-^H2D^b-^, referred to as HLA-C*05^+^ KO cells). Cells were labelled with differential concentrations of CFSE to allow for cellular discrimination as follows: KO cells (0.5μM of CFSE), WT cells (5μM CFSE) and HLA-C*05^+^ KO (5μM CFSE). Cells were mixed and ∼ 5 x 10^6^ cells per population were injected intravenously into either WT or KIR2DL1-expressing recipient mice. Splenocytes were isolated from recipient mice after 20 hours and cells were analysed by flow cytometry using antibodies for HLA-C (B1.23.2) and H2K^b^ (48-5958-82). A small sample of the injection mix was kept and analysed by flow cytometry for reference. Survival of both HLA-C*05^+^ KO and KO cells was calculated relative to the WT cells in each experiment using the injection mix as reference. Relative survival of HLA-C*05^+^ KO compared to survival of KO cells was plotted.

### Isolation of thymic epithelial cells (TECs)

TECs were isolated as previously described (Jain and Gray, 2014). In brief, thymus lobes were collected from 6-7 week old mice, were cleaned from fat and connective tissue and the capsule was removed under a dissection microscope. Cleaned tissue was digested in PBS, 0.4 Wunsch Units/ml Liberase^TM^ (Sigma-Aldrich) and 300μg/ml DNaseI (VWR) pre-warmed to 37°C. Tissue was carefully triturated in a step-wise manner. First, after 5 min incubation in 37°C water bath, the thymus lobes were carefully pipetted up and down 10-15 times using a cut P1000 pipette tip. Tissue fragments were left to settle down for 5 min, the cell suspension transferred into a collection tube containing R10 and cells pelleted. Cells were re-suspended in FACS buffer containing 50μg/ml DNaseI to prevent cell clumping. Trituration of the remaining tissue fragments was repeated a total of four rounds; with pipette tip openings gradually getting smaller and the last round using an uncut P10 tip. Cells were pooled and counted. An aliquot was taken for flow cytometric staining as a pre-TEC isolation sample. CD45^+^ cells were depleted using CD45 MicroBeads (Miltenyi Biotech) according to manufacturer’s instructions. Cells were filtered sequentially through 70μm and 40μm (Corning) cell strainers with 1.8mg/ml DNaseI added in order to prevent blocking of the MACS columns. The CD45^−^ cell fraction eluted from the LS columns (Miltenyi Biotech) represented TECs. Cells were counted and flow cytometric surface marker staining was performed as described above.

### Lentiviral transduction and *in vitro* co-culture assays

The HLA-C*05 DNA construct (used to make transgenic mice) and a similarly designed HLA-C*07 control DNA construct were sub-cloned into the lentiviral pHRsinUbEm expression plasmid (a gift from J.M. Boname/ P.J. Lehner, University of Cambridge) as described previously (Kaur et al., 2017). To confirm that the presence of the H2-K^b^ α3 domain in the HLA-C construct did not hamper recognition of the HLA-C molecule by KIR2DL1, two additional constructs, C*05 (α3-H) and C*07 (α3-H) constructs, were made. In the C*05 (α3-H) and C*07 (α3-H) DNA constructs, the α3 domain and hence the exons 4-8 of the HLA-C molecule were taken from the respective HLA-C genes. The lentiviral HLA-C expression plasmids were co-transfected with the vesicular stomatitis virus-G envelope plasmid pMD2.G (Addgene) and packaging plasmid psPAX2 (Addgene), containing HIV-1 Gag and Rev, into HEK 293T cells to package lentiviral particles. Viral titres were determined by serial dilution and transduction of HEK 293T cells. 721.221 cells were transduced with the packaged lentivirus at a multiplicity of infection of 20, in the presence of polybrene (Santa Cruz Biotechnology), added at a final concentration of 8 µg/ml. Cells were harvested 72 hours post transduction and HLA-C expression determined by staining with anti-HLA (W6/32) (Biolegend) antibody. The HLA-C transduced 721.221 cells were then co-cultured with YT-KIR2DL1 cells at a ratio of 0.5:1 T:E for 6 hours and IFN*γ* production by the YT-KIR2DL1 cells was assessed by staining with antibodies for IFN*γ* (XMG1.2) and KIR2DL1 (HP-MA4). In addition, controls where the empty-vector transduced 721.221 cells or non-transduced 721.221 cells were co-cultured with YT-KIR2DL1 cells, or where YT-KIR2DL1 cells were cultured with PMA and Calcium Ionomycin were included.

### Isolation and culture of trophoblast cells

Trophoblast cells were isolated from implantation sites from age-matched pregnant female mice at gd12.5 as previously described (Rodrigues-Duarte et al., 2018). All placentas from the litter from a mouse were pooled and minced in RPMI. For every 6-8 placentas, 25ml digestion buffer containing HEPES (20 mM) (Gibco), sodium bicarbonate (0.35 g/L) (Gibco), collagenase type I (1mg/ml) (Sigma-Aldrich), DNAse I (20 µg/ml) in RPMI supplemented with Glutamax was used and incubated at 37°C for 1 hour. Cells were centrifuged at 500g for 5 min, and the pellet was re-suspended in 4 ml of 25% Percoll and layered on top of 4 ml of 40% Percoll, followed by centrifugation at 850g for 20 min at 4°C without brake. Trophoblast cells were removed from the interface, washed with PBS, incubated with ACK lysis buffer followed by another wash in PBS. Cells were plated at 1×10^5^ cells/well in 96-well U-bottom plates and cultured in RPMI supplemented with Glutamax, 10% heat inactivated FCS, sodium pyruvate, penicillin/streptomycin, L-glutamine and 50µM 2-mercaptoethanol for 4 days, with media replenished after 2 days. On day 3, half the cells were stimulated with LPS (10µg/ml) and IFN-γ (500U/ml), and the other half were cultured without stimulus. Cells were then harvested using Accutase cell dissociation reagent (Fisher Scientific) after 24 hours and stained as per the flow cytometric staining protocol.

### ELISA

Mouse VEGF R1/Flt-1 (R&D systems), PIGF-2 (R&D systems) and Endoglin/CD105 (R&D systems) were measured in plasma samples collected from pregnant mice on gd18.5. For collection of plasma, murine blood was collected in EDTA-coated tubes followed by centrifugation at 2000g for 15 min at room temperature. Supernatants (plasma) were collected, aliquoted and stored at −80°C. Urine samples were also collected from pregnant female mice at gd18.5 (just prior to culling the mice for measurement of fetal weights), and aliquoted and stored at −80°C. Albumin and creatinine concentrations in the urine were determined by an Albumin ELISA (Bethyl Laboratories) and a Creatinine colorimetric assay (Cayman Chemical), respectively. The ratio of urinary albumin to creatinine was calculated and considered as a measure of proteinuria.

### RNA isolation, cDNA preperation and Q-PCR

Total RNA was isolated from cells using the RNeasy isolation kit (Qiagen), and cDNA prepared using the Quantitech Reverse Transcription kit (Qiagen). SYBR-green based quantitative PCR assays were deigned and optimised for *HLA-C* and *Hprt*. Samples collected from tissue such as dissected implantation sites from pregnant mice or organs were cut into small pieces and immersed in RNAlater (Qiagen) as per manufacturer’s instructions, prior to processing for RNA isolation. The sequences of the primers used for the assays are provided in the Table S3. Standard curves for each of the assays were performed using serial dilutions of cDNA and amplification efficiencies were determined. Relative expression was expressed as 2^−ΔCt^, where ΔCt is the difference of the cycle threshold between the transcript of the gene of interest and the reference gene transcript.

### RNA *in situ* hybridization (RNA-ISH)

RNA-ISH was performed on mouse implantation site tissues isolated from pregnant mice on gd9.5. Implantation sites were embedded in OCT and snap frozen in isopentane and dry ice. Serial 10µm coronal sections were cut on a cryostat, kept at −20°C to dry for an hour and then stored at −80°C. Sections were hybridized with mRNA probes specific for the *Ccl1*, *Gzme/d/g* and *Anxa2* genes (ACD). Following probe hybridization and amplification, mRNA was detected using the RNAscope 2.5 HD Duplex Assay (ACD), as per the manufacturer’s instructions. For quantification of stainings, tissue from four independent implantation sites were used per mouse group with multiple sections from the same region of the tissue used from each implantation site. Image analysis was completed by the Pharma Services group at Indica Labs (Albuquerque, NM, USA). Slides were scanned at 40X magnification on the Aperio Scanscope Turbo AT2 Slide Scanner (Leica Biosystems) and digital images imported into HALO^®^ Image Analysis Platform (Indica Labs, Albuquerque, NM USA). Collaboration between the two groups occurred through the use of a cloud-based HALO^®^ Link Image Management system. Image analysis was based on teal-red duplex RNAScope^®^ images counterstained with hematoxylin (Sigma-Aldrich) for nuclei detection. HALO^®^ Link was utilized to import sample metadata and for researcher annotation of samples for regions of interest (ROIs). Annotations were used to identify individual tissue sections on separate layers of analysis. Multiple HALO^®^ algorithms were utilized for analysis: ISH v3.3.9 (RNA probe identification) and Multiplex IHC v2.1.1 (cytoplasmic and nuclear staining localisation). The ISH algorithm was optimized to detect positive intensity of individual “spot-like” probes of *Anxa2* and *Ccl1*. *Anxa2* displayed the expected “spot-like” staining pattern of RNAScope^®^ while *Ccl1* displayed a stronger “spot-like” staining pattern than whole cell or nuclear staining, and both of these were accurately identified using ISH. In addition to the ISH algorithm we also utilized our Multiplex IHC algorithm because the staining pattern for *Gzme/d/g* was not a typical probe-like RNA stain. Therefore, the Multiplex IHC algorithm more accurately quantified this probe. The Multiplex IHC algorithm was optimized to detect positivity within the nuclear compartment of *Gzme/d/g*. The ISH module calculated the Probe Copies/cell for *Anxa2* and *Ccl1*, and the Multiplex IHC module calculated the overall % positive cells for *Gzme/d/g*.

### Blood pressure measurement in mice

Systolic blood pressure was measured in mice using a non-invasive blood pressure analyser, BP-2000 Blood Pressure Analysis System (Bioseb), that uses tail-cuff transmission photoplethysmography. The restraint platform was kept at a temperature of 37°C. Baseline blood pressure measurements for each mouse were taken prior to using the mice for timed-matings. Baseline measurements required recording of valid measurements for at least 3 days for each mouse with a minimum of 4 valid readings on each day of measurement. During the course of gestation, measurements were taken every alternate day from gd2.5 until gd14.5 and then every day from gd15.5 until gd18.5. Post-partum measurements were taken daily from day 1 post-partum until 6-7 days post-partum. Inclusion of measurements during pregnancy and post-partum for each mouse required recording of a minimum of 4 valid readings on each day of measurement. Collected data were analysed with the BP-2000 Analysis Software (Bioseb) and the change in blood pressure for each mouse was normalised to its baseline.

### Perfusion of uteroplacental vasculature for Micro-CT imaging and vascular segmentation analysis

The method for injection of the silicon rubber injection compound, Microfil (Flow Tech) into the uteroplacental vasculature of mice for micro-CT imaging has been previously described (Rennie et al., 2014). In brief, pregnant mice at gd10.5 were given terminal anaesthesia using pentobarbitone and intracardiac perfusion was performed using a Heparin solution (100 IU/ml) in PBS. A catheter was placed in the descending thoracic aorta and used to clear the lower body vasculature of blood and perfused with heparinised PBS. The contrast agent, Microfil was then gently infused by hand on a slow flow rate until the blue colour of the contrast agent was visible through the capillary bed and sufficiently vented through the right atrium. The inferior vena cava was then immediately tied off and the vasculature was kept pressurised using a syringe that contained the contrast agent to sustain vessel inflation, until the Microfil polymerised. The uterus was then removed, immersed in formalin (Sigma-Aldrich) overnight and the implantation sites were mounted in 1% agarose for micro-CT imaging. Micro-CT imaging was performed on Skyscan 1172 (Bruker) at 40kV, 250μA, 750ms exposure time and a pixel size of 4.73μm, and the three-dimensional (3D) images were reconstructed using Nrecon (Micro Photonics). As the uteroplacental vasculature was difficult to segment automatically due to the unique nature of the tightly coiled spiral arteries, the micro-CT datasets were manually segmented slice by slice using the seg3D software (“Seg3D”). Each acquired image stack was first downscaled by 50% (to typical dimensions 900×900×1000) and 3D median filtering was applied (radius 2 voxels, isotropic). Each voxel of the resulting stack was classified as background, vessel or transitional using multi-Otsu thresholding. With the aid of these pre-assigned non-overlapping classes, assisted manual segmentation was performed using seg3D, segmenting the volume into spiral arteries, radial arteries, maternal canals, uterine arteries, and background. The first step of the automated analysis was to translate the voxel-based information in the segmented image stacks into a graph structure that is better suited for quantification algorithms. The uterine, spiral, radial and maternal canal vessels were combined into a single class and skeletonisation was performed using homotopic thinning (Lee et al., 1994). The skeletonised network was then organised into segments, where a segment denotes the ordered chain of vertices that run between two adjacent junctions, or a junction and a terminal. Spurious segments that may result from noisy segmentation were detected using a custom routine and removed from the network (Lee et al., 2007). All subnetworks (a collection of segments that are connected to each other) were detected and the smallest subnetworks below the threshold (< 30 segments) were removed. Vessel class labelling from the segmentation was automatically transferred to the segments. Because the skeletonisation process places nodes at discrete voxel centres, the total number of nodes in a given network depended on the total length of the vessels in it. To allow inter-sample comparisons, we chose to normalise the number of total nodes in each network to 10,000 by resampling along each segment. The distance between each adjacent pair of nodes was kept approximately constant, subject to the constraint of a discrete number of voxels along a segment. At each node, the local radius of the vessel segment was estimated using Rayburst algorithm (Rodriguez et al., 2006). Furthermore, to refine the voxel-based quantification of radius, a sequence of synthetic vessel-like image stacks with known radii were created and nonlinear regression was used to calibrate the Rayburst estimates into sub-voxel quantities. Using the Rayburst kernel, the location of each skeleton node was also adjusted towards the true vessel centreline on a sub-voxel basis, to reduce the staircase artefact. Vessel morphological quantifications were performed either on a nodal basis for radii calculations, or segmental basis for calculation of length and volume, for vessels > 10μm in radius and length. For segmental quantification, the summed segment length and volume were calculated using a piecewise linear approximation. Flow conductance of each segment was calculated as the reciprocal of Poiseuille resistance, under the assumption that the flow will remain laminar. Given the typical inlet vessel dimensions, this assumption was regarded as appropriate. However, the calculation of total spiral arteries conductance requires further assumptions about boundary conditions. For this, the terminal vessels of the spiral subnetwork were detected and their proximity to the radial or canal vessels were evaluated, thereby classifying every terminal as being radial- or canal-adjacent. All radial-connected spiral arteries were assumed as flow inlets and all canal-connected arteries, as outlets. In general, the number of inlets and outlets are not consistent between different samples. We calculated the total spiral network resistance by imposing a pressure gradient between the inlets and outlets of 1kPa and calculating the total outflow using a network-Poiseuille formulation, as 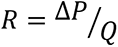. Due to the linearity of the flow model, the resistance remains constant regardless of the actual pressure gradient applied. For analysis, flow conductance is quoted, which are reciprocal of the resistance (the greater the conductance, the easier the fluid to flow).

### Isolation of cells from implantation sites and FACS sorting for scRNA-seq studies

Isolation of cells from the implantation sites were done with enzymatic digestion using Liberase^TM^ as previously described (Collins et al., 2009). In brief, uterine horns were collected in cold HBSS containing Ca^2+^ and Mg^2+^ (ThermoFisher) from pregnant mice and cleared of mesometrial fat. The tissue was collected from the entire mesometrial side of the implantation site leaving behind the developing fetus (Edwards et al., 2014; Pang et al., 2014). All implantation sites from a litter were pooled and processed together. The tissue was finely minced, incubated in HBSS solution containing 0.28 Wunsch Units Liberase^TM^ and 30μg/ml DNase for 30 minutes at 37°C with agitation, washed with Ca^2+/^Mg^2+^-free PBS containing 5mM EDTA (ThermoFisher) and centrifuged. Cells were incubated in Ca^2+/^Mg^2+^-free PBS containing 5mM EDTA for 15 min at 37°C, followed by filtering through a 70μm cell strainer, red blood cell lysis and washing in PBS. Cells were then filtered again through a 70μm cell strainer and counted.

For isolation of total unsorted cells from the implantation site, half of the isolated cells were taken and depleted of dead cells by labelling with the Dead cell removal microbeads (Miltenyi Biotech) as per the manufacturer’s instructions. The viable cell fraction eluted from the MS columns (Miltenyi Biotech) was counted and processed for droplet based scRNA-seq (below).

For isolation of uNK cells, half of the isolated cells were surface stained using the flow cytometric staining protocol for antibodies against CD45 (30-F11), CD3 (145-2C11), CD19 (eBio1D3), TCRb (H57-597), NKp46 (29A1.4), CD49a (Ha31/8), CD49b (DX5), KIR2DL1 (HP-MA4), CD122 (TM-b1) along with either 7AAD or a Live/Dead viability dye. For SMART-Seq2 processing, single cells were FACS sorted into 96 U-well plates into either trNK (NCR1^+^CD3ε^−^TCRb^−^CD19^−^CD45^+^CD122^+^CD49a^+^) or cNK (NCR1^+^CD3ε^−^TCRb^−^CD19^−^CD45^+^CD122^+^CD49b^+^) cells. Single cells were directly sorted into wells containing 5μl of pre-chilled TCL buffer (Qiagen) containing 1% 2-mercaptoethanol as previously described (Trombetta et al., 2014). Each plate consisted of a negative control with no cells and a positive control with 15 cells sorted in one well. 96-well plates containing the sorted cells were then immediately centrifuged and frozen directly on dry ice prior to storage at −80°C. For droplet based scRNA-seq, total uNK cells (NCR1^+^CD3ε^−^TCRb^−^CD19^−^CD45^+^CD122^+^) were sorted into Eppendorf tubes containing PBS with 0.04% BSA and counted and processed as described below.

### Full length scRNA-Seq

SMART-Seq2 scRNA-seq libraries were prepared from trNK and cNK cells sorted in 96 U-well plates as previously described (Picelli et al., 2014; Trombetta et al., 2014) with modifications in the protocol as previously described (Ding et al., 2020). Briefly, RNA was purified from single cell lysates using 2.2x RNAClean XP beads (Beckman Coulter), eluted in 4µl of master mix containing 1µl of 10 mM dNTPs (ThermoFisher), 0.1µl of 100 µM 3’ oligodT RT primer, 0.1 µl of 40 U/µl RNase Inhibitor (Clontech Takara) and 2.8 µl of 1 M Trehalose (Life Sciences Advanced Technologies), followed by incubation at 72°C for 3 min and placing on ice. The remaining reverse transcription mix, containing 2µl of Maxima RT buffer (ThermoFisher), 0.1 µl of 100 µM TSO oligo, 0.25 µl of 40 U/µl RNase Inhibitor, 0.1 µl of 200 U/µl Maxima H Minus Reverse Transcriptase (ThermoFisher), 3.45 µl of 1 M Trehalose and 0.1 µl of 1 M MgCl_2_ (Sigma-Aldrich) was added to each well and incubated for 50°C for 90 min followed by 85°C for 5 min. The cDNA was then amplified using 12.5 µl of KAPA HiFi HotStart ReadyMix (KAPA Biosystems) and 0.05 µl of 100 µM ISPCR primer and incubated at 98°C for 3 min followed by 23 cycles of 98°C for 15 sec, 67°C for 20 sec, 72°C for 6 min, and a final extension at 72°C for 5 min. PCR products were purified using 0.7x AMPure XP SPRI beads (Beckman Coulter) and eluted in 20 µl of TE buffer (Qiagen). DNA quantification was done using the Qubit™ dsDNA HS Assay Kit (ThermoFisher) and DNA fragment size was assessed using the High Sensitivity DNA BioAnalyzer kit (Agilent). cDNA concentrations were normalised and 0.125 ng of each cDNA was used in a quarter volume of a Nextera XT DNA library preparation kit (Illumina), following pooling of equal volumes from each well and purification with 0.6x AMPure XP SPRI beads. Final libraries were again assessed for DNA concentration using the Qubit assay and fragment size using the Bioanalyzer. All SMART-Seq2 Libraries were sequenced using a NextSeq 500/550 High Output v2 kit 75 cycles (Illumina) on a Nextseq 500 (Illumina).

### Droplet based scRNA-Seq

For droplet based scRNA-seq, 7,000 cells were loaded onto the Chromium Single Cell 3’ Chip A (10x Genomics) and processed through the chromium controller to generate Gel beads in Emulsion. RNA-seq libraries were prepared using the Chromium Single Cell 3’ Library & Gel Bead Kit v2 (10x Genomics) as per the manufacturer’s instructions. Libraries were sequenced on an Illumina Hi-Seq.

### Statistical analysis

Data were analysed using Prism 8 or RStudio. Statistical details of experiments including sample size can be found in the figure legends. To statistically estimate the effect of genotype on fetal or placental weight, a linear mixed-effects model was used with transgenic genotype mating combination as a fixed effect and clustering of observations within a litter as a random effect. Similarly, for estimation of the effect of genotype on spiral artery diameter and segment length, a linear mixed-effects model was used with transgenic genotype mating combination as a fixed effect and clustering of observations from a specific implantation site as well as clustering of observations within a litter from a mouse as random effects. As the segment length data followed a log-normal distribution, the mixed effects model for assessing effect on segment length was run on log-transformed length values. For comparison of two-group datasets with non-Gaussian distribution, the non-parametric Mann-Whitney test was used. Values were expressed as mean ± S.E.M. P values lower than 0.05 were considered statistically significant. For all the statistical analysis: *p < 0.05, **p < 0.01, ***p < 0.001, ****p < 0.0001.

### scRNA-Seq data preprocessing

For full length scRNA-Seq SMART-Seq2 data, raw sequencing data was demultiplexed to fastq files using bcl2fastq (v.2.17.1.14) and aligned to a custom reference using RSEM (v.1.2.8); the custom reference included the mm10 genome and the sequence for the human genes *KIR2DL1* and *HLA-C*05* (as described in the transgenic mice generation above), and was generated using *rsem-prepare-reference*. Quality control was assessed and summaries generated using STAR. We used RSEM (v1.2.8) to quantify gene counts and TPM (we converted TPM to TP100K after initial quality control filtering, described below).

For droplet-based scRNA-Seq data, raw sequencing data was demultiplexed to fastq files with *cellranger mkfastq* (version 2.1.0, 10x Genomics) and reads were aligned and count matrices were generated with *cellranger count* (version 2.0.1). A custom reference was generated to include the *HLA-C*05* and *KIR2DL1* transgenic construct sequences into the mm10 reference using *cellranger mkref* (version 2.1.0). The alignment of *HLA-C*05* sequence was limited to the human DNA sequence region to minimise the risk of misalignment with the mouse *H-2K^b^* gene.

### scRNA-Seq quality control

For full length scRNA-seq, lower-quality cells were removed if they met any of the following criteria: (**1**) log_10_(counts)<5, (**2**) number of expressed genes <1,000 or > 7,000, or (**3**) average housekeeping gene (**Table S1**) expression (TPM) > 1. In addition, cells that met any of the following criteria were also removed (to ensure cells meet FACS expectations): (**1**) Cells with HLA-C*05 expression <2 in C*05/KIR x WT mating group; (**2**) Cells with HLA-C*05 expression > 2 in WT x C*05 mating group, (**3**) Cells with KIR2DL1 expression < 2 in KIR x C*05 and C*05/KIR x WT mating groups; (**4**) Cells with KIR2DL1 expression > 2 in WT x C*05 mating group; (**5**) Cells with Cd3e expression > 3; (6) In sorted cNK cells, cNK cells with Itga1 expression > 1. Genes expressed in three or fewer cells were removed. Final expression values were normalized to TP100K and log transformed.

For droplet-based scRNA-seq, the feature-barcode matrix cellranger count was used, but with *force-cells=6000* to ensure all droplets containing cells are included. Low quality cells were removed by filtering any cells with either (**1**) less than 500 genes, (**2**) less than 1000 UMIs, or (**3**) more than 10% of UMIs mapped to mitochondrial genes. Genes expressed in three or fewer cells were removed. The final expression values were normalized to a total UMI per cell of 1×10^4^ and log transformed.

We used Scrublet (v.0.2.1), with *expected_doublet_rate = 0.06*, to predict doublets (Wolock et al., 2019). In the NK cell sorted droplet-based data we removed one cluster with high doublet probability as well as one cluster of non-NK cells, and then re-analyzed the remaining data. In the unsorted droplet-based data we excluded from visualization and analysis cells from two clusters with high doublet scores, but did not reanalyze the data following exclusion.

### Dimensionality reduction, clustering and visualization

In both full length and droplet-based scRNA-seq, human genes were removed from the expression matrix prior to dimensionality reduction. We did not batch-correct by mouse because mice and genotype were evenly distributed across clusters (**Fig. S4C,F**,**I,L,S7E,** adjusted rand index between sample and cluster = 0.49% for full length NK data, 0.88% for droplet-based NK data, and 1.21% for the droplet-based unsorted data).

For both the full-length and droplet-based scRNA-seq data, highly variable genes were selected using *FindVariableGenes* (default parameters) in Seurat (Butler et al., 2018) [v.2.1]. For the full-length data, the expression of each gene was centered and scaled to have a mean of zero and a standard deviation of one; for droplet-based data, the expression of each gene was centered to have a mean of zero.

In both full length and droplet-based data sets, initial dimensionality reduction was performed by principal component analysis (PCA) using *RunPCA* in Seurat (v.2.1) (Butler et al., 2018), and Louvain graph-based clustering was performed using *FindClusters* (based on the top 15 PCs, with clustering resolutions of 0.8, 0.4, and 0.6 for the full-length NK, droplet-based NK, and unsorted droplet-based data, respectively; other parameters, including *k*=30 for *k*-NN, were set to the default). Cell profiles were visualized using a Uniform Manifold Approximation and Projection (UMAP) (McInnes et al., 2018) embedding of the top 15 PCs, with *min_dist=0.5*, *spread=1.0*, *number of neighbors = 15*, and the Euclidean distance metric.

The statistical significance of genotype proportions within each cluster was tested using a Dirichlet-multinomial regression (Smillie et al., 2019). Mouse batch effects were assessed within each cluster by calculating the rand index using the Clues package in R.

### Differentially expressed genes, annotating cell subsets, and building gene signatures

Subset-specific marker genes (all data sets) and cNK/trNK-specific (full-length data) differentially expressed genes were identified by a Mann-Whitney U test using the *FindAllMarkers* function in Seurat (v.2.1), with *test.use = ‘wilcox’*, *logfc.threshold* = *log(1.5)* and *min.pct = 0.2*; the Benjamini-Hochberg False Discovery Rate (FDR) was calculated using *p.adjust* in R (*method = ‘Benjamini & Hochberg’*). Cell subsets were manually annotated using cell subset marker genes and expression of relevant published signatures (Aran et al., 2017; Nelson et al., 2016; Pavlicev et al., 2017; Tirosh et al., 2016; Tsang et al., 2017; Vento-Tormo et al., 2018; Wallrapp et al., 2017). For the two NK cell scRNA-seq data sets (full-length and droplet-based) and some cell subsets from the unsorted droplet-based data, novel cell subsets are described by their expressed marker genes.

Within each subset, we tested for differentially expressed genes between IUGR and CTR1, IUGR and CTR2, and CTR1 and CTR2 using: (1) a pseudobulk differential expression analysis and (2) a mixed-effects Poisson regression model. In both approaches, genes were only tested for differential expression if they were expressed in at least 10% of cells within one of the tested conditions for a given cell subset. For the pseudobulk approach, we summed counts per mouse within a subset and then performed differential expression analysis using DESeq2 (v.1.26, design formula: *gene ∼ condition*) (Love et al., 2014). We used the *fdrtools* (v.1.2.16) package (Strimmer, 2008) in R to calculate FDR based on the empirical null, with the DESeq2 Wald statistic as input. For the mixed-effects Poisson regression model (Subramanian et al., 2020), we used the *glmer* function from the *lme4* package (v.1.1-26) (Bates et al., 2015) in R with *family = poisson()* and the following design formula: *gene ∼ condition + offset(log(total_UMI/mean_UMI)) + (1|mouseID)*. We calculated FDR using *p.adjust* in R (*method = ‘Benjamini & Hochberg’*). IUGR-unique expression changes were defined as genes that were differentially expressed in the IUGR *vs*. CTR1 (FDR < 0.05), but not differentially expressed in CTR1 *vs*. CTR2 (FDR < 0.05). Volcano plots of differentially expressed genes were made using the R package, EnchancedVolcano (v.1.4.0) (Blighe et al., 2019).

Gene signatures for the cNK and trNK cells and for NK cell subsets were defined as the union of the top 25 differentially expressed genes (sorted by p-value) and the top 25 differentially expressed genes sorted by log fold change. The proliferation signature was previously reported (Wallrapp et al., 2017). Gene signatures were scored in each cell using *AddModuleScore* in Seurat (Butler et al., 2018) (v.2.1 for scoring proliferation and v.3.2.2 for scoring NK cell subsets).

### Topic modeling

Topic models were fit to the droplet-based sorted uNK UMI counts using the *FitGoM()* function from the CountClust (v.1.6.1) R package (Bielecki et al., 2018; Dey et al., 2017; Xu et al., 2019). To select the number of topics, *K*, models are fit for each of a range of *K* (K=4:22 in increments of two) with a tolerance parameter (tol) of tol=0.1 and tol=0.01. The Bayesian Information Criterion (BIC), estimated likelihood, and Akaike Information Criterion (AIC) were computed using *compGoM()* for each model. The final value of K (K=16) was chosen by considering the BIC curve (seeking a K for which the BIC was either minimizing or decreasing slowly) and the biological relevance of the identified gene programs. The final value of tol=0.01 was chosen primarily by the biological relevance of the identified gene programs. Top scoring genes per topic were defined using the *extractTopFeatures* function.

Topics significantly changed in IUGR were selected by examining four criteria: (**1**) the sign of the KS statistic; (**2**) the Benjamini Hochberg FDR following the KS-test; (**3**) the fold change direction and magnitude of the mean; and (**4**) the fold change direction and magnitude of the median. The *ks.test* and *p.adjust* (with *method = ‘Benjamini & Hochberg’*) functions in R were used to calculate the FDR for IUGR *vs.* CTR1 and IUGR *vs.* CTR2 under the KS test. The KS-statistic was calculated with the *ks.test* function in R as *cumsum(ifelse(order(w) <= n.x, 1/n.x, −1/n.y))*, where *n.x = length(x)*, *n.y = length(y)*, and *w = c(x,y)*. The final statistic was reported as the maximum or minimum of the above equation, whichever had the larger absolute magnitude. We examined the above four criteria using 100% of cells as well as cells with cell weights in the top 75%, 50%, and 25% for each topic; we report metrics based on cells with weights in the top 75% as this retained the most number of cells without a skew towards cells with very low weights in a topic.

EnrichR (Chen et al., 2013; Kuleshov et al., 2016) and Ingenuity Pathway Analysis (Qiagen) were used to calculate pathway enrichment. Enrichment of canonical pathways, diseases and functions was performed in IPA using gene lists of IUGR-unique expression changes (defined as genes that were differentially expressed in IUGR *vs*. CTR1 but not differentially expressed in CTR1 *vs.* CTR2 within each subset; above); the background was the intersection of genes that were in IPA with the genes that were expressed in the cell subset in at least 10% of the cells from either condition.

### Cell-cell interactions

Mouse genes were mapped to their human genes orthologs using the Human Phenotype Ontology(http://www.informatics.jax.org/downloads/reports/HMD_HumanPhenotype.rpt). CellPhoneDB (Vento-Tormo et al., 2018) was then used to identify individual significant ligand-receptor interactions between subsets in the unsorted droplet-based scRNA-Seq data with *cellphonedb method statistical_analysis meta.txt counts.txt --result-precision=3 --iterations=1000*, where *meta.txt* is cell subset assignments, *counts.txt* is the normalized counts data for either the WT x C*05 or KIR x C*05 matings, *result-precision=3* indicated the number of decimal points for the reported p-value, and *iterations* was the number of iterations for the statistical test.

The number of significant interactions was summed for each subset pair, and interactions were visualized using Cytoscape v3.8.2 (Shannon et al., 2003). Only subset pairs with at least 50 interactions were connected on the Cytoscape map, and line thickness corresponded to the number of significant interactions.

### Data and code availability

Raw data, gene expression matrices and the code used to generate results presented in this paper will be made available upon publication

“Seg3D”, S.C.a.I.I. “Seg3D” Volumetric Image Segmentation and Visualization. Scientific Computing and Imaging Institute (SCI).

